# ARID1A-BAF coordinates ZIC2 genomic occupancy for epithelial to mesenchymal transition in cranial neural crest lineage commitment

**DOI:** 10.1101/2024.04.03.587869

**Authors:** Samantha M. Barnada, Aida Giner de Gracia, Cruz Morenilla-Palao, María Teresa López-Cascales, Chiara Scopa, Francis J. Waltrich, Harald M.M. Mikkers, Maria Elena Cicardi, Jonathan Karlin, Davide Trotti, Kevin A. Peterson, Samantha A. Brugmann, Gijs W. E. Santen, Steven B. McMahon, Eloísa Herrera, Marco Trizzino

## Abstract

The BAF chromatin remodeler regulates lineage commitment including cranial neural crest cell (CNCC) specification. Variants in BAF subunits cause Coffin-Siris Syndrome (CSS), a congenital disorder characterized by coarse craniofacial features and intellectual disability. Approximately 50% of CSS patients carry variants in one of the mutually exclusive BAF subunits, *ARID1A/ARID1B*. While *Arid1a* deletion in mouse neural crest causes severe craniofacial phenotypes, little is known about the role of ARID1A in CNCC specification. Using CSS patient-derived *ARID1A*^+/-^ iPSCs to model CNCC specification, we discovered *ARID1A*-haploinsufficiency impairs epithelial to mesenchymal transition (EMT), a process necessary for CNCC delamination and migration from the neural tube. Furthermore, wild-type ARID1A-BAF regulates enhancers associated with EMT genes. ARID1A-BAF binding at these enhancers is impaired in heterozygotes while binding at promoters is unaffected. At the sequence level, these EMT enhancers contain binding motifs for ZIC2, and ZIC2 binding at these sites is ARID1A-dependent. When excluded from EMT enhancers, ZIC2 relocates to neuronal enhancers, triggering aberrant neuronal gene activation. In mice, deletion of *Zic2* impairs NCC delamination, while *ZIC2* overexpression in chick embryos at pre-migratory neural crest stages elicits ectopic delamination from the neural tube. These findings reveal a novel ARID1A-ZIC2 axis essential for EMT and CNCC delamination.

## Introduction

The mammalian SWI/SNF complex, referred to as BRG1/BRM associated factor (BAF), is an ATP-dependent chromatin remodeler^1^. BAF plays global roles in lineage specification^2–4^, pluripotency^5^, tumorigenesis^6^, and basic cellular processes^7–9^ by modulating chromatin accessibility and interacting with transcription factors^10^ to impact gene regulation. BAF comprises three main modules: 1) the ATPase module that hydrolyzes ATP for catalytic activity to alter nucleosome positioning, 2) the ARP module which aids in ATPase module functions, and 3) the core module required for complex assembly, stabilization, and DNA-binding^11,12^. The three BAF subtypes, canonical BAF, polybromo-associated BAF (PBAF), and non-canonical (GLTSCR1/1L-containing) BAF (GBAF)^5,13^ include all the main modules in varying configurations which contain interchangeable subunits that confer context-dependent functions^14–16^. Canonical BAF (hereafter “BAF”) is the only subtype that contains the large, mutually exclusive <u>A</u>T-rich interaction <u>d</u>omain <u>1A</u> and <u>1B</u> (ARID1A/ARID1B) subunits in its core module^11,12,15^.

ARID1A and ARID1B are necessary for BAF complex formation, stability, and DNA-binding^12,15,16^. *De novo* alterations in *ARID1A* can occur at all stages from germline mutations to somatic mosaicism in embryonic development^17,18^ and adult tissue^6,19^. In cancer, ARID1A is reported to have context-dependent oncogenic^20,21^ and/or tumor suppressive functions^22–25^ as variants in *ARID1A* occur in ∼10% of all tumors^6,26^, and ∼50% of specific cancer types^24,27^. In a developmental setting, heterozygous loss-of-function variants in *ARID1A* (as well as mutations in other BAF subunits) are causative of Coffin-Siris Syndrome (CSS)^28,29^ due to haploinsufficiency. CSS is a rare developmental disorder characterized by systemic congenital anomalies including distal limb phenotypes, intellectual disability, and coarse craniofacial features^30–32^. These dysmorphic craniofacial features including a depressed nasal bridge, short nose, averted nares, broad philtrum, and wide mouth^32^, result from impaired craniofacial development.

The process of forming the craniofacial skeleton relies on complex spatiotemporal regulation and patterning and includes derivatives of all three germ layers and neural crest cells^33^. Neural crest cells (NCCs) are a transient cell population that arise between 3-4 weeks of human embryonic development during neurulation^34^. Post-gastrulation, a portion of the ectoderm begins to differentiate into neuroectoderm, forming the neural plate that is flanked by the neural plate border^34^. During neurulation, the neural plate invaginates and separates from the bordering non-neural ectoderm at the neural plate border to form the neural tube^34^. At this neural plate border zone, NCCs undergo epithelial to mesenchymal transition (EMT), delaminate, and migrate to populate their terminal sites in the developing embryo^35–37^.

Distinctively, NCCs are multipotent and will form characteristic ectodermal derivatives, such as peripheral neurons, as well as form cell types that are typically mesodermal-derived such as cardiac tissue and craniofacial bone and cartilage^38^. Cranial neural crest cell-derived bone and cartilage form the frontal, nasal, zygomatic, maxillary, mandibular, and dentary bones, as well as the bones of the inner ear and the hyoid^37,39^. Moreover, malformations of these bones underlie many of the craniofacial anomalies seen in CSS patients including a depressed nasal bridge, short nose, averted nares, broad philtrum, and wide mouth^32^. Similarly, *Arid1a* conditional knock-out models in neural crest of developing mouse embryos^40^ recapitulate these craniofacial phenotypes characteristic of CSS. Together, the clinical manifestations of *ARID1A*-haploinsufficient individuals and the craniofacial anomalies in conditional *Arid1a^-/-^* mouse models demonstrate the indispensable role of ARID1A-BAF in neural crest formation and function. However, the molecular pathways regulated by ARID1A during craniofacial development and cranial neural crest specification remain poorly understood.

Herein, we sought to uncover the role of ARID1A-BAF in the specification and migration of CNCCs in the context of craniofacial development. Our data suggests that ARID1A-BAF is required to activate gene networks associated with EMT. Using induced pluripotent stem cells (iPSCs) derived from *ARID1A*^+/-^ CSS patients, we discovered impaired ARID1A correlates with the upregulation of neuronal networks at the expense of EMT networks. Motifs matching the binding site for the ZIC2 transcription factor were highly enriched at the ARID1A-bound genes linked to EMT, and *in vitro* and *in vivo* experiments suggest ARID1A-mediated induction of EMT occurs via ZIC2. These findings highlight a novel axis involving an ARID1A-ZIC2 interaction required for CNCC EMT and suggests a pathogenic mechanism that may underlie the ARID1A-associated CSS craniofacial phenotypes.

## Results

### Control and Coffin-Siris patient-derived iPSCs are pluripotent

We leveraged control and *ARID1A* variant-containing human-derived iPSCs, to investigate the role of ARID1A in the BAF chromatin remodeling complex during cranial neural crest specification and craniofacial development. To generate these iPSCs, skin fibroblasts were collected from two Coffin-Siris Syndrome patients each harboring distinct, previously reported heterozygous, *de novo*, nonsense mutations in *ARID1A*^17^ (OMIM: 614607). Patient 1 (PT1) cells originated from a teenage male harboring a *ARID1A*:c.1113delG;p.Gln372Serfs*19 variant and Patient 2 (PT2) cells originated from a female harboring a *ARID1A*:c.3679G>T;p.Glu1227* variant (Fig. 1A). Both variants were classified as pathogenic based on the criteria set forth by the American College of Medical Genetics and Genomics^41^. A subsequent *ARID1A*- wildtype iPSC line was used as a control (CTRL1). While full mutations in *ARID1A* lead to a severe CSS subtype with multiple congenital anomalies and reduced survival, mosaic variants are often observed in individuals with a more moderate CSS phenotype^42^. The *ARID1A*-haploinsufficient patient fibroblasts (PT1 and PT2) were reprogrammed into iPSCs (Supplementary Fig. 1A) and due to the associated mosaicism, it was possible to isolate *ARID1A* wild-type iPSCs (CTRL2) in addition to the *ARID1A*-haploinsufficient iPSCs from PT2 for an isogenic study system. Genotypes of the resulting iPSC lines were verified via deep sequencing.

**Figure 1.**
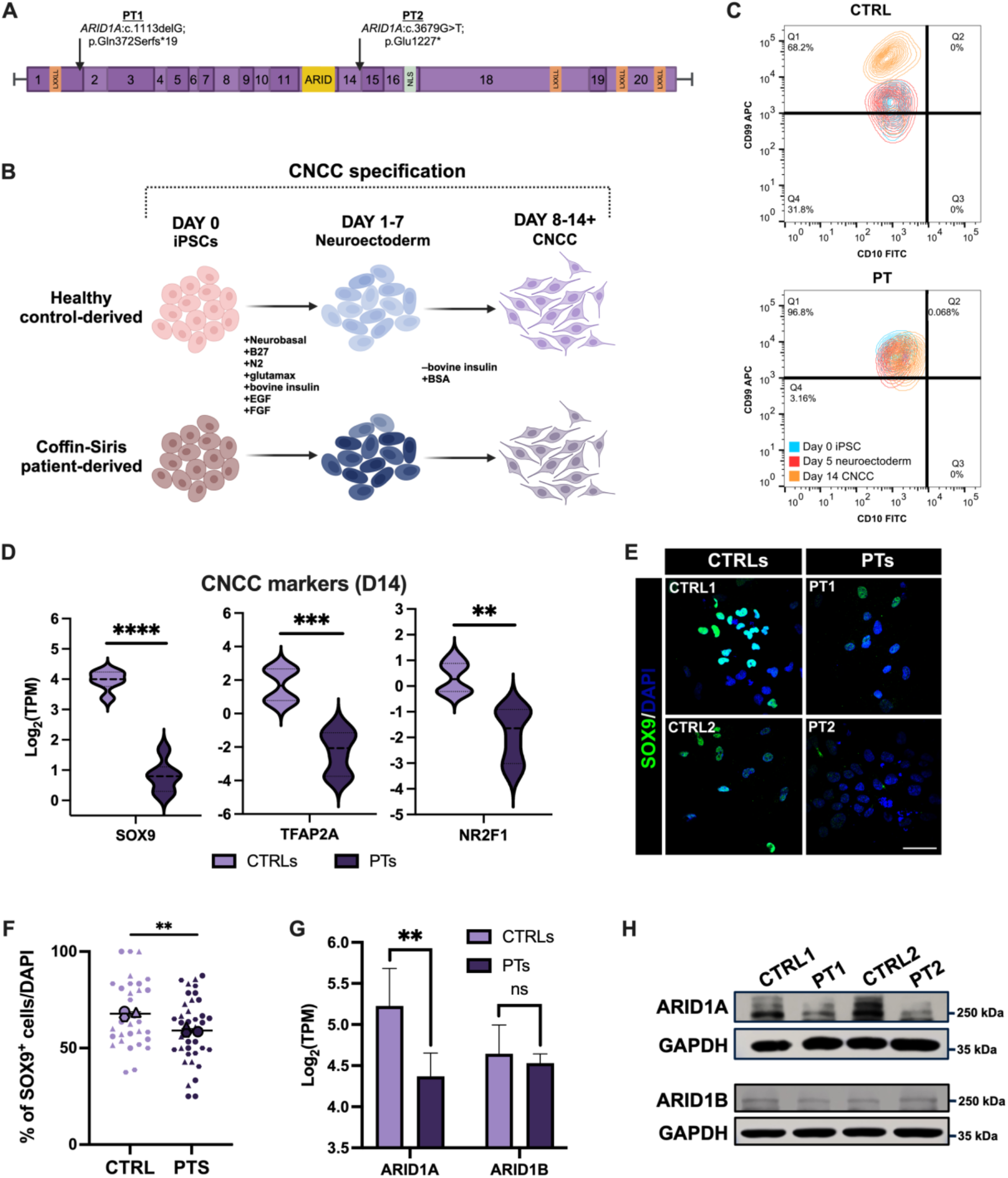
ARID1A-haploinsufficient cells fail to specify as CNCCs. (A) Schematic of the *ARID1A* gene (NM_ 006015.6) highlighting the two nonsense mutations in each patient cell line. Each number represents a corresponding exon in *ARID1A*. LXXLL = protein-interaction domain; ARID = AT-rich interaction domain or DNA-binding domain; NLS = nuclear localization signal. Made with BioRender.com. (B) Graphical illustration of 2D *in vitro* CNCC specification. EGF = epidermal growth factor; FGF = fibroblast growth factor; BSA = bovine serum albumin. Made with BioRender.com. (C) PT lines at day 14 (D14) fail to show a distinctive increase in the CD99 surface marker representative of successful CNCC specification. All populations (D0 iPSC, D5 neuroectoderm, D14+ CNCC) are CD10^+^ with a slight increase in the CD10 surface marker at D14 CNCC which is characteristic of preparation of a differentiation trajectory towards the mesenchymal lineage. (D) Violin plots displaying log_2_(TPM) of CNCC markers SOX9, TFAP2A, and NR2F1 at D14 between CTRLs and PTs. A two-tailed unpaired t-test was performed and p < 0.05 was considered significant; SOX9: ****p < 0.0001, TFAP2A: ***p = 0.0001, and NR2F1: **p = 0.0012. (E) Representative images and (F) quantification of a SOX9 immunofluorescence performed at D14 in CTRL and PT lines. (E) DAPI staining on nuclei in blue. Merged images shown taken at 60X magnification; scale bar = 50µm. (F) SuperPlot quantification of percentage of SOX9^+^ cells per DAPI. n = 3 represented by a distinctive shape with each small data point representing a captured image. The larger data points correspond to the average values of each replicate. A two-tailed unpaired t-test performed on the average values between CTRLs and PTs; p < 0.05 was considered significant; **p = 0.0032. (G) Log_2_(TPM) bar plot and (H) Immunoblot of ARID1A and ARID1B at D10 of CNCC specification between CTRLs and PTs. Reduction of ARID1A transcript and protein levels correspond to the ARID1A-haploinsufficient genotype while ARID1B levels are consistent across CTRLs and PTs. A two-tailed unpaired t-test was performed and p < 0.05 was considered significant; ARID1A: **p = 0.0029, ARID1B: ns p = 0.4592.

Previous studies from our lab^3^ and others^43,44^ have shown that ARID1A-containing BAF is present in pluripotent stem cells (Supplementary Fig. 1B). Additionally, GBAF (non-canonical BAF) is an alternate embryonic stem cell-specific configuration of BAF which lacks ARID subunits and instead incorporates GLTSCR1/GLTSCR1L and BRD9^5,13^. Notably, GBAF is also present in embryonic stem cells where it is implicated in regulating pluripotency^5^. Consistent with the lack of ARID subunits in GBAF, the *ARID1A*- haploinsufficient PT iPSCs display regular iPSC morphology, including small cells making up distinct, circular, compact colonies with well-defined colony borders (Supplementary Fig. 1C), and broadly express OCT4, SOX2, and NANOG mRNA and protein (Supplementary Fig. 1D-F). RNA-seq of iPSCs identified only 121 differentially expressed genes between CTRL and PT cells (p < 0.05; FDR < 5%; log_2_(fold change)+/-1; Supplementary File 1). Notably, genes involved in pluripotency regulation were not differentially expressed (Supplementary File 1). These data show that CSS patient iPSCs maintained pluripotency and *ARID1A*-haploinsufficiency did not impact pluripotent gene expression in this cell type.

### CSS cranial neural crest cells exhibit migratory defects

As premature loss of pluripotency did not appear to be the pathological mechanism of ARID1A- CSS, we next investigated the role of ARID1A in cranial neural crest specification. CTRL and CSS PT iPSCs were specified to CNCCs as per previously published protocols^45–47^ (Fig. 1B). RNA-seq analysis comparing CTRL iPSCs (Supplementary File 2) and derived CNCCs (Supplementary File 3) revealed a transcriptional signature in which CNCC markers were highly expressed while pluripotency genes were downregulated (Supplemental Fig. 1G), supporting successful CNCC specification (Supplementary Fig. 1H).

Previous work from our group demonstrated a bimodal switch between ARID1A and ARID1B in the BAF complex during the CNCC specification process^3^ (Supplementary Fig 1B). At the onset of neuroectoderm formation, ARID1A protein expression is downregulated and nearly undetectable while ARID1B protein expression increases^3^ (Supplementary Fig. 1B). ARID1B remains highly expressed throughout the duration of the neuroectodermal stage (days 1-7) but it is then downregulated during the onset of EMT and CNCC formation. At this stage, ARID1A protein levels are re-upregulated. This latter switch occurs at day 8-9 of our CNCC specification protocol, and ARID1A is retained in the BAF complex as migratory CNCCs are formed^3^ (Supplementary Fig. 1B). Notably, this time-point immediately precedes EMT and delamination of the CNCCs as they acquire a migratory phenotype.

To investigate the impact *ARID1A*-haploinsufficiency has on the formation of CNCCs, we harvested specified CNCCs on day 14. At this stage, CTRL cells displayed characteristic single-cell CNCC morphology, while PT cells adopted a more elongated and clustered cellular morphology (Supplementary Fig 1I). By day 14, PT cells failed to form the distinct CD99^+^ CNCC population which is characteristic of successful specification (Fig. 1C; Supplementary Fig. 2) as observed by flow cytometry. Contrary to results observed in iPSCs, RNA-seq performed in CTRL and PT CNCCs at day 14 revealed over 3,800 differentially expressed genes (p < 0.05; FDR < 5%; log_2_(fold change) +/-1; Supplementary File 4). Notably, genes associated with CNCC specification and migration (i.e. *SOX9*, *TFAP2A*, and *NR2F1*), were significantly decreased in the PT cells relative to CTRL CNCCs (Fig. 1D). Immunostaining at day 14 validated RNA-seq data showing a reduced number of SOX9^+^ PT cells relative to CTRLs (Fig. 1E & 1F).

As ARID1A expression returns by day 8-9 of our in vitro CNCC specification program^3^, we chose day 10 for comprehensive genomic studies, focused on the role of ARID1A in CNCC formation. First, we wanted to elucidate if other ARID proteins (i.e. ARID1B or the PBAF member ARID2) could compensate for *ARID1A*-haploinsufficiency. As expected, ARID1A protein levels were significantly diminished in the PT lines at day 10 of CNCC formation (Fig. 1G & 1H). However, neither ARID1B nor ARID2 were upregulated at the transcript (Fig. 1G; Supplementary Fig. 1J) or protein level (Fig 1H) at the same timepoint, suggesting other BAF configurations could not compensate for the partial loss of ARID1A during CNCC formation. Together, these data implicated ARID1A-BAF as a key regulator of CNCC formation.

### ARID1A haploinsufficiency results in decreased expression of EMT genes and increased expression of neuronal genes

In addition to the 3,800 genes differentially expressed at the end of the CNCC specification protocol (day-14, Supplementary File 4), RNA-seq analyses performed at day-10 of CNCC specification (i.e. at the onset of ARID1A re-upregulation) identified 652 differentially expressed genes in PT lines compared to CTRLs (Fig. 2A; Supplementary File 5). Of these 652 genes, 349 were downregulated and 303 were upregulated in PT lines. Downregulated genes were significantly enriched in pathways associated with epithelial to mesenchymal transition (Fig. 2B) while the upregulated genes were significantly enriched in neuronal pathways (Fig. 2C & 2D). Notably, both neural progenitors and NCCs are derived from the neuroectoderm (Fig. 2E). We previously demonstrated that this developmental precursor was regulated by a specific version of BAF incorporating ARID1B rather than ARID1A^3^. However, the *ARID1A*- haploinsufficient lines failed to upregulate EMT and CNCC genes while aberrantly upregulating neuronal genes. Thus, the partial loss of ARID1A in CSS patient cells appeared to impair formation of the multipotent CNCC lineage from neuroectoderm, triggering a default differentiation to the neuronal lineage (Fig. 2E). Further, we examined transcript levels of neuronal markers at the conclusion of the CNCC specification process (day 14). Neuronal markers *MAP2*, NeuN (*RBFOX3*), and *DCX* were significantly upregulated in PT lines compared to CTRLs (Fig. 2F). Moreover, immunofluorescence for MAP2 and NeuN revealed a massive increase in expression of both markers in the PT lines (Fig. 2G & H). These data suggest that the *ARID1A*- haploinsufficient lines were unable to commit towards the CNCC lineage, while aberrantly upregulating a mix of neural progenitor and neuronal markers.

**Figure 2.**
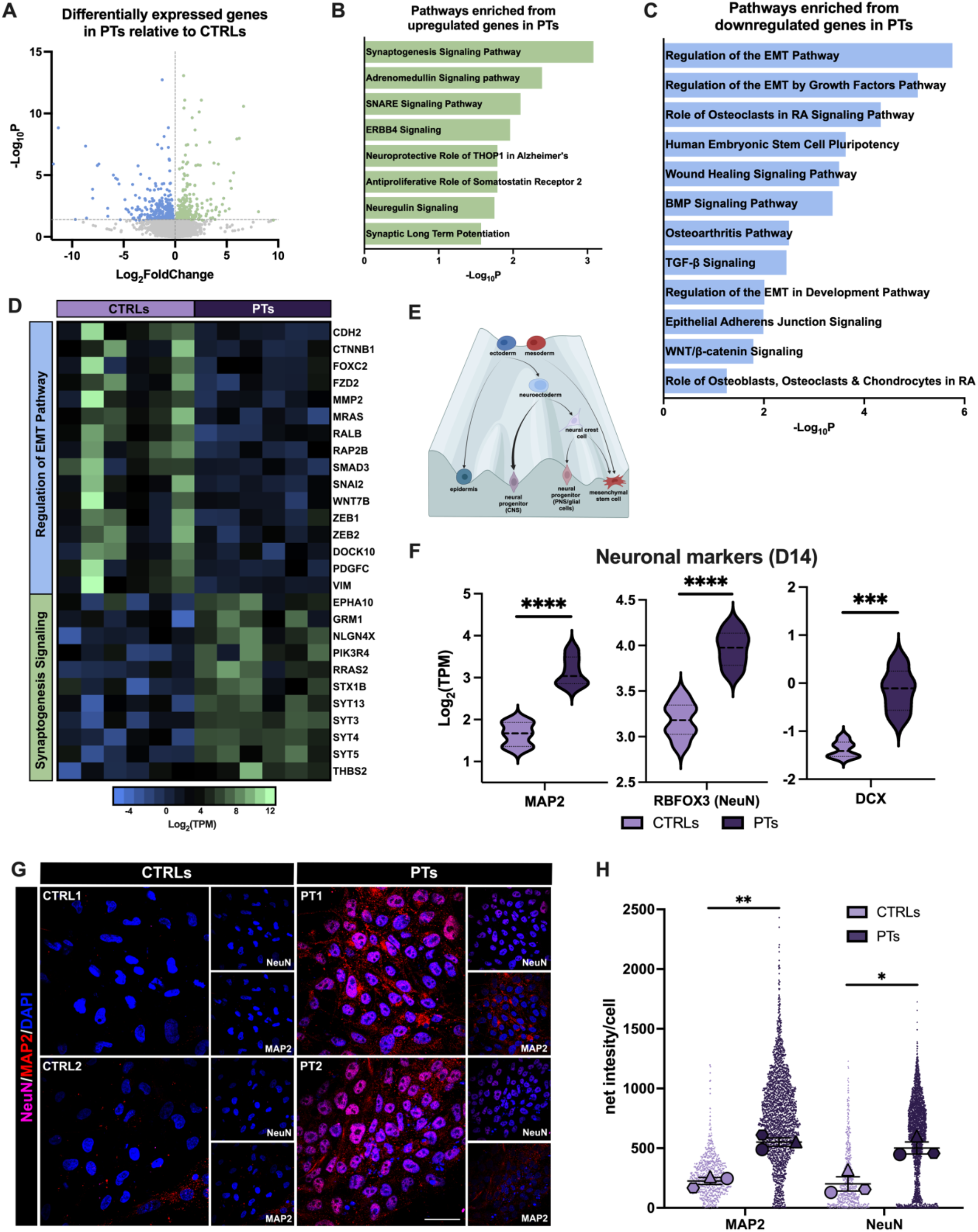
PT cell lines do not undergo EMT and default to a neuronal lineage commitment. (A) Volcano plot of differentially expressed genes in PTs relative to CTRLs at D10 of CNCC specification as determined by DESeq2. Green points represent aberrantly upregulated genes while blue points represent aberrantly downregulated genes. n = 652. (B) Neuronal pathways are enriched from differentially upregulated genes in PTs at D10 of CNCC specification and the respective -log_10_p-value as determine by IPA. (C) Pathways enriched from differentially downregulated genes in PTs at D10 of CNCC specification and the respective -log_10_p-value as determine by IPA. Identified downregulated pathways are essential for successful CNCC specification including epithelial to mesenchymal transition (EMT), BMP, TGF-β, and Wnt signaling. RA = rheumatoid arthritis. (D) Heatmap of genes involved in the most significant upregulated and downregulated pathways presented in (B) and (C) between CTRLs and PTs. Genes involved in the Regulation of EMT Pathway were downregulated while there was an increase in expression of genes involved in Synaptogenesis Signaling Pathway in PT lines. CTRL and PT columns represent 3 individual replicates from 3 separate CNCC specifications for CTRL1/CTRL2 and PT1/PT2 respectively. (E) Adapted schematic of Waddington’s epigenetic landscape depicting restrictive cell fate commitment in the context of neural crest formation. The mesoderm gives rise to mesenchymal stem cells, and the ectoderm differentiates into the epidermis and neuroectoderm. Neuroectodermal fate defaults to the neural progenitor lineage of the central nervous system (CNS) represented by a thick arrow when neural crest fate cannot be specified. Neuroectoderm also gives rise to multipotent neural crest cells which will differentiate to mesenchymal stem cells for formation of various derivatives including the craniofacial bones and cartilage and neural progenitors and glia of the peripheral nervous system (PNS). Made with BioRender.com. (F) Violin plots displaying log_2_(TPM) of neuronal markers MAP2, RBFOX3 (NeuN), and DCX at D14 between CTRLs and PTs. A two-tailed unpaired t-test was performed and p < 0.05 was considered significant; MAP2: ****p < 0.0001, RBFOX3 (NeuN): ****p < 0.0001, and DCX: ***p = 0.0001. (G) Representative images and (H) quantification of MAP2 and NeuN immunofluorescence performed at D14 in CTRL and PT lines. (G) DAPI staining on nuclei in blue. Images taken at 60X magnification; scale bar = 50µm. (H) SuperPlot quantification of net intensity of MAP2 and NeuN per cell. n = 3 represented by a distinctive shape with each small data point representing a captured image. The larger data points correspond to the average values of each replicate. A two-tailed unpaired t-test performed on the average values between CTRLs and PTs; p < 0.05 was considered significant; MAP2 **p = 0.0021 NeuN *p = 0.0184.

### Functional EMT is impaired in CNCCs in ARID1A haploinsufficiency

CNCCs are a multipotent population with the potential to differentiate into both ectodermal and mesenchymal derivatives^38^ (Fig. 2E). To migrate and differentiate into the terminal derivatives, CNCCs must undergo epithelial to mesenchymal transition (EMT) allowing for delamination and migration from the neural tube^35–37^. Key mechanisms involved in developmental EMT include regulation of cadherin switching whereby E-cadherin is downregulated, and N-cadherin is upregulated^48–50^. This switch is accompanied by increased expression of essential mesenchymal factors vimentin, SNAIL, SLUG, and the TWIST, and ZEB transcription factor families^48^.

Notably, we found that markers defining epithelial/neuroectodermal cells, including E-cadherin (*CDH1*) and *EPCAM* and pre-migratory CNCCs (*FOXD3*), were all upregulated in the *ARID1A*- haploinsufficient lines compared to CTRL CNCCs (day 14; Fig. 3A). Conversely, mesenchymal and migratory CNCC markers, such as N-cadherin (*CDH2*), *TWIST1*, and vimentin (*VIM*) were markedly downregulated in the *ARID1A*-haploinsufficient lines compared to CTRL CNCCs (Fig. 3B). Immunostaining for the epithelial marker EpCAM (Fig. 3C & 3D), and the mesenchymal marker vimentin (Fig. 3E & 3F), further supported the transcriptomic data, strongly indicating that the PT lines remained in a neuroepithelial state, while the CTRLs successfully underwent EMT.

**Figure 3.**
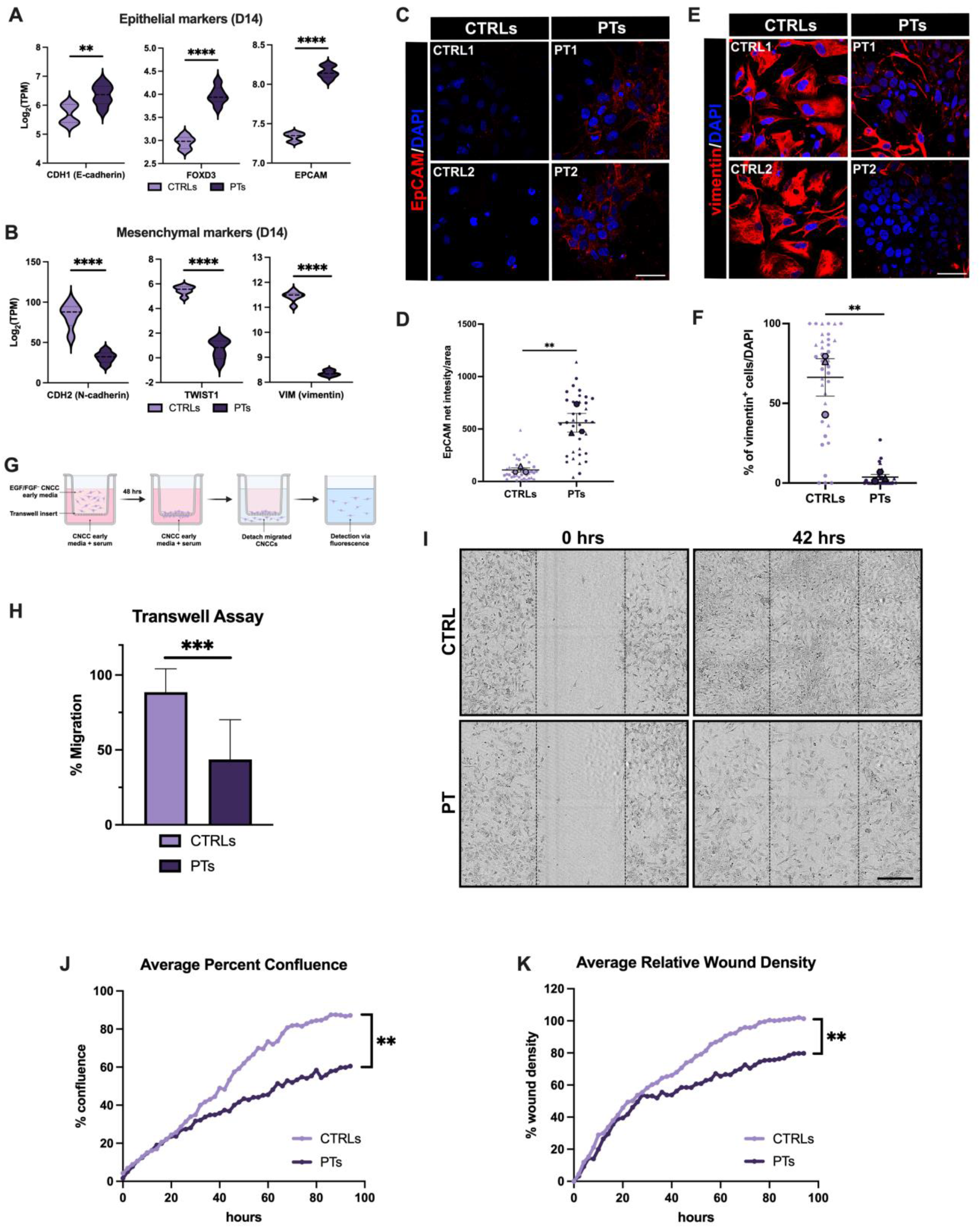
Impaired EMT compromises the specification of ARID1A-haploinsufficient cells into migratory CNCCs. (A) Violin plots displaying log_2_(TPM) of epithelial/pre-migratory CNCC markers CDH1 (E-cadherin), FOXD3, and EPCAM at D14 between CTRLs and PTs. A two-tailed unpaired t-test was performed and p < 0.05 was considered significant; CDH1 (E-cadherin): **p = 0.0079, FOXD3: ****p < 0.0001, and EPCAM: ****p < 0.0001. (B) Violin plots displaying log_2_(TPM) of mesenchymal/migratory CNCC markers CDH2 (N-cadherin), TWIST1, and VIM (vimentin) at D14 between CTRLs and PTs. A two-tailed unpaired t-test was performed and p < 0.05 was considered significant; CDH2 (N-cadherin): ****p < 0.0001, TWIST1: ****p < 0.0001, and VIM (vimentin): ****p < 0.0001. (C) Representative images and (D) quantification of an EpCAM immunofluorescence performed at D14 of CNCC specification in CTRL and PT lines. (C) DAPI staining on nuclei in blue. Merged images shown taken at 60X magnification; scale bar = 50µm. (D) SuperPlot quantification of EpCAM net intensity per area. n = 3 represented by a distinctive shape with each small data point representing a captured image. The larger data points correspond to the average values of each replicate. A two-tailed unpaired t-test performed on the average values between CTRLs and PTs; p < 0.05 was considered significant; **p = 0.0078. (E) Representative images and (F) quantification of a vimentin immunofluorescence performed at D14 of CNCC specification in CTRL and PT lines. (E) DAPI staining on nuclei in blue. Merged images taken at 60X magnification; scale bar = 50µm. (F) SuperPlot quantification of percentage of vimentin-positive cells per DAPI. n = 3 represented by a distinctive shape with each small data point representing a captured image. The larger data points correspond to the average values of each replicate. A two-tailed unpaired t-test performed on the average values between CTRLs and PTs; p < 0.05 was considered significant; **p = 0.0062. (G) Graphical illustration of a transwell assay. Made with BioRender.com. (H) Bar plot depicting average percent migration of CTRL and PT cells through a transwell membrane. A two-tailed unpaired t-test performed between CTRLs and PTs; p < 0.05 was considered significant; ***p = 0.0002. (I) Representative brightfield images of a scratch wound assay (Incucyte) at 0 hours (hrs) and 42 hrs post-scratch between CTRL and PT lines. 42 hrs represents the earliest time point of a visually closed wound in CTRL cells. Scale bar = 300µm, 10X magnification. (J) Average percent confluence of the wound area and (K) average relative wound density over 96 hours between CTRLs and PTs from the scratch wound assay performed in (I). A two-tailed unpaired t-test performed between CTRLs and PTs; p < 0.05 was considered significant; percent confluence: **p = 0.0014, percent wound density: **p = 0.0077.

To functionally assess the migratory potential of CTRL and PT lines *in vitro*, we performed transwell (Fig. 3G-H) and scratch wound assays (Fig. 3I-K). After CNCCs were specified, growth factors were removed from the media to allow for an accurate assessment of cell migration. PT cells had approximately 50% reduced migration than CTRLs through the transwell membrane (Fig. 3H). Similarly, the CTRL scratch wound was visually confluent while the PT line exhibited minimal migration (Fig. 3I). A significant decrease in average percent confluence in the scratch wound as well as relative wound density was observed in PT lines compared to CTRLs (Fig. 3J & 3K). Collectively, these data suggest that EMT was affected in PT lines as they failed to acquire a functional migratory phenotype.

### ARID1A-BAF regulates accessibility at EMT enhancers

To investigate the mechanism underlying the impaired EMT in the PT lines, we performed chromatin immunoprecipitation followed by sequencing (ChIP-seq) for ARID1A to assess alterations in binding at day-10 (i.e. ARID1A re-upregulation onset). Simultaneously, we performed an assay for transposase-accessible chromatin with sequencing (ATAC-seq) to interrogate chromatin accessibility at the same time point. We detected 21,724 shared ARID1A ChIP peaks across PT and CTRL cells (Supplementary Fig. 3A), 3,398 new ARID1A peaks gained in PT lines, and 19,401 ARID1A peaks lost in PT lines (Fig. 4A & 4B). Similarly, there were ∼46,065 conserved ATAC-seq peaks between PTs and CTRLs (Supplementary Fig. 3B), 9,163 peaks exclusive to PT cells, and 20,216 peaks exclusive to CTRL cells (Supplementary Fig. 3C & 3D). Consistent with *ARID1A* loss being heterozygous, average profiles indicate that both ARID1A-binding and chromatin accessibility at the ∼20,000 sites were nearly half of the peak height of the CTRL levels (Fig. 4B and Supplemental Fig. 3E). Moreover, the loss of ARID1A binding and accessibility were correlated (Fig. 4C).

**Figure 4.**
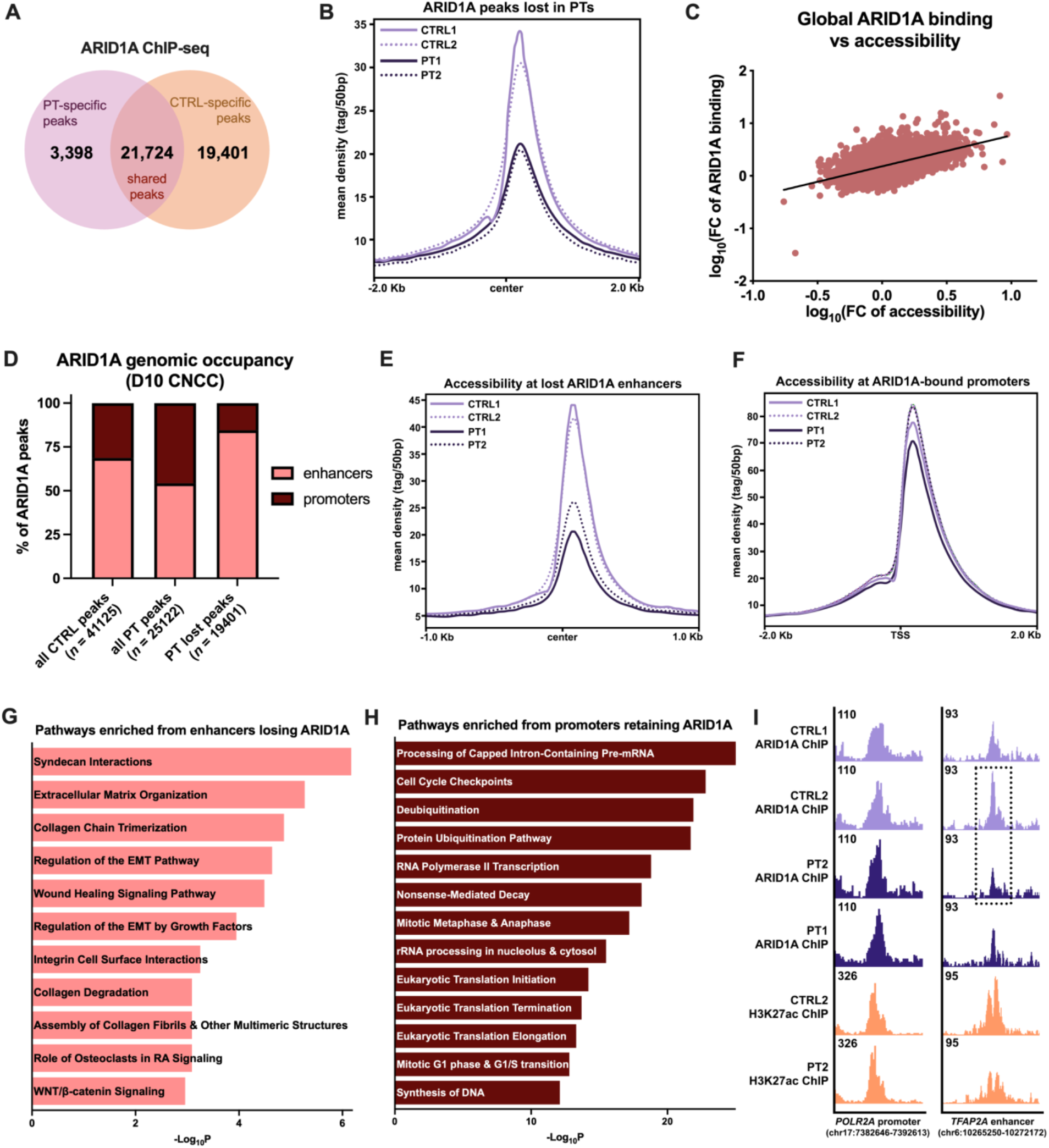
Chromatin dynamics and ARID1A genomic occupancy in CTRL and PT cell lines. (A) Venn diagram displaying the number of PT-specific peaks (ARID1A peaks present only in PTs or “gained” in PTs; 3,398), CTRL-specific peaks (ARID1A peaks present only in CTRLs or “lost” in PTs; 19,401), and shared ARID1A peaks between CTRLs and PTs (conserved ARID1A binding; 21,724) via ARID1A ChIP-seq performed at D10 of CNCC specification. (B) Average profile of ARID1A peaks lost in PTs at D10 (n = 19,401). Center represents the average overlapping global binding of ARID1A at CTRL-specific peaks. (C) Scatter plot and linear regression of the log_10_(fold-change) of ARID1A peak coverage versus the log_10_(fold-change) of ATAC peak coverage displaying a significant Pearson correlation (r = 0.4940, two-tailed ****p <0.0001) between global ARID1A binding and global accessibility. (D) Stacked bar plot depicting the percentage of ARID1A peaks enriched at enhancers (>1kb from the closest transcription start site or TSS) and promoters (<1kb from the closest TSS) in all CTRL ARID1A peaks (including peaks shared with PT lines), all PT ARID1A peaks (including peaks shared with CTRL lines), and all ARID1A peaks that are lost in the PTs (PT lost peaks). ARID1A peaks in PTs are enriched at promoter regions relative to CTRLs; 54% at enhancers and 46% at promoters in PTs versus 69% at enhancers and 31% at promoters in CTRLs. Approximately 85% of lost ARID1A peaks in the PT lines were lost at enhancer regions. (E) Average profile of ATAC peaks at enhancer regions that lose ARID1A binding and accessibility in the PTs at D10 of CNCC specification (n = 3,203). Center represents the average overlapping lost PT ATAC/ARID1A peaks at enhancer regions across the genome. (F) Average profile of ATAC peaks at promoter regions that retain ARID1A binding and accessibility in the PTs at D10 of CNCC specification (n = 11,231). TSS represents the average overlapping retained ATAC peaks at transcription start sites throughout the genome. (G) Pathways enriched from genes that lose ARID1A binding at their enhancers in PTs at D10 of CNCC specification and the respective -log_10_p-value as determined by IPA. Pathways include those necessary for migration, extracellular matrix dynamics, and EMT. (H) Pathways enriched from genes that retain ARID1A binding at their promoters in PT lines at D10 of CNCC specification and the respective -log_10_p-value as determined by IPA. Pathways include those necessary for basic cellular functions and survival including DNA synthesis, transcription, translation, and cell cycle. (I) Integrative Genomics Viewer example of a retained ARID1A peak in the PTs at the promoter of POLR2A, encoding the main subunit of RNA-polymerase II, and a lost ARID1A peak at an enhancer for TFAP2A, encoding an essential CNCC marker. Representative promoter and enhancer regions marked by H3K27ac. Dotted black box highlights the decrease in ARID1A binding in the isogenic system.

Analyzing the genomic occupancy of ARID1A, approximately 70% of the CTRL ARID1A peaks were at enhancer regions (>1kb from the closest transcription start site) and 30% at promoters (<1kb from the closest transcription start site; Fig 4D). Conversely, ARID1A peaks in the PT lines were enriched equally between enhancers and promoters (Fig. 4D). The majority of the ARID1A peaks lost in the PTs were at enhancer regions (85% at enhancers and 15% at promoters), while most of the ARID1A peaks at promoters were retained (Fig 4D). This suggests that BAF binding at promoter regions may be essential, while BAF binding at enhancers may be dispensable (Fig. 4D). Consistent with this, accessibility at ARID1A-bound enhancers was significantly diminished in PT lines (Fig. 4E), while accessibility at ARID1A-bound promoter regions was largely retained (Fig. 4F). Pathway analysis on differentially expressed genes associated with enhancers that lost ARID1A binding revealed enrichment for EMT, wound healing, extracellular and collagen dynamics, and integrin interactions (Fig. 4G). However, pathway analysis on genes associated with ARID1A-regulated promoters revealed enrichment for basic cellular processes including cell cycle, transcription, translation, and DNA synthesis (Fig. 4H). For example, the promoter of *POLR2A*, encoding the main subunit of RNA-polymerase II, harbors conserved ARID1A occupancy, while a *TFAP2A* enhancer displayed loss of both ARID1A binding and H3K27ac signal (Fig. 4I). These data suggested that ARID1A-BAF regulates EMT by opening the chromatin at EMT-associated enhancers, and the function of these enhancers is impaired in *ARID1A*-haploinsufficiency.

### ARID1A-BAF regulates the accessibility at enhancers of EMT genes to coordinate ZIC2 binding

To predict potential interactions between ARID1A and transcription factors in the regulation of EMT enhancers during CNCC specification, we performed a motif analysis on the ARID1A-bound enhancers and promoters. Promoters that retained ARID1A binding harbored conventional promoter motifs (Supplementary Fig. 4A), while the EMT enhancers that lose ARID1A binding in PT cells revealed motifs for CNCC lineage-specific transcription factors^4,45,51–57^ (Fig. 5A). Strikingly, nearly 40% of these enhancers harbored a ZIC family motif. ZIC1 is not expressed at this developmental stage^58,59^, while ZIC2 and ZIC3 are both highly expressed in CNCCs^60–64^. Reportedly, *ZIC2* mutations cause holoprosencephaly type 5 in humans^65^, a developmental syndrome which shares many phenotypic features^66^ with Coffin-Siris^30–32^. Therefore, we investigated ZIC2 as a potential interactor of ARID1A in EMT regulation.

**Figure 5.**
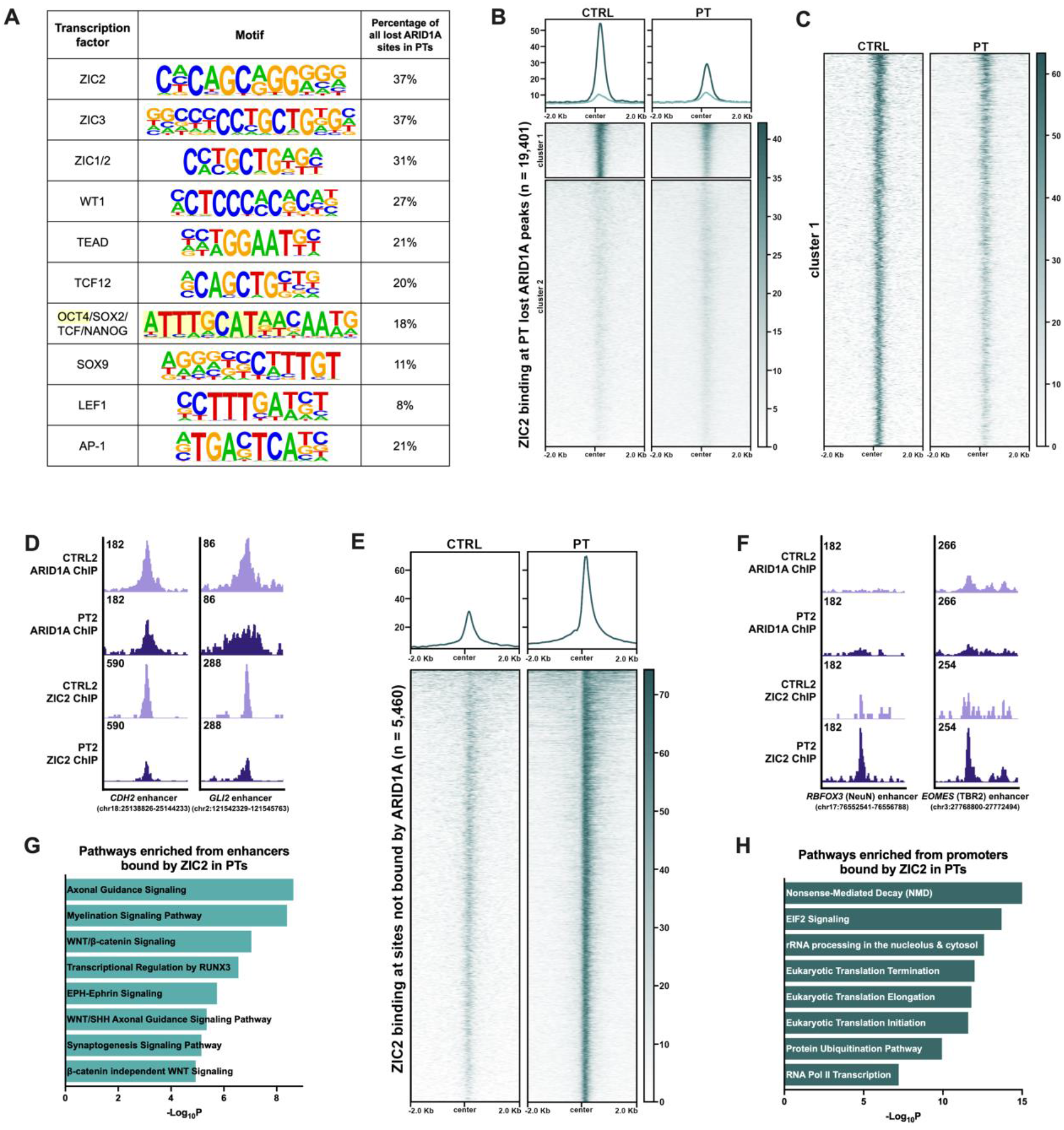
ZIC2 regulates CNCC and neuronal identity by activating accessible regulatory networks. (A) Table of enriched motifs at ARID1A-bound regions that are lost in PTs identified by HOMER. The presence of the motifs at each lost region was quantified via FIMO (MEME Suite). (B) Clustered heatmap of ZIC2 binding at regions that lose ARID1A binding in the PT lines at D10 of CNCC specification (CTRL-specific; n = 19,401). (C) Expanded heatmap of cluster 1 from (B). (D) Integrative Genomics Viewer example of lost ARID1A and ZIC2 peaks in PT2 relative to the isogenic CTRL2 at ARID1A-target enhancers at D10 of CNCC specification. *CDH2* and *GLI2* are necessary for mesenchymal identity and migration of CNCCs respectively. (E) Heatmap of ZIC2 binding at regions that are not bound by ARID1A displaying a genomic relocation of ZIC2 in *ARID1A*-haploinsufficient conditions at D10 of CNCC specification (n = 5,460). (F) Integrative Genomics Viewer example of gained ZIC2 binding in PT2 relative to the isogenic CTRL2 at sites not bound by ARID1A at D10 of CNCC specification. *RBFOX3* (NeuN) and *EOMES* (TBR2) are neuronal markers. (G) Pathways enriched from enhancers that are bound by ZIC2 in PT2 at D10 of CNCC specification and the respective -log_10_p-value as determined by IPA. Pathways include those involved in neuronal signaling and processes. (H) Pathways enriched from promoters that are bound by ZIC2 in PT2 at D10 of CNCC specification and the respective -log_10_p-value as determined by IPA. Pathways include those necessary for basic cellular functions and survival including transcription, translation, and nonsense-mediated decay.

To determine whether ZIC2 requires ARID1A to bind to EMT enhancers, we performed a ChIP-seq for ZIC2 in our isogenic CTRL and PT lines. In *ARID1A*-haploinsufficient conditions we observed attenuation of ZIC2 binding at EMT enhancers normally regulated by ARID1A (Fig. 5B & 5C; Supplementary Fig. 4B). Similar to ARID1A, ZIC2 also preferentially bound enhancers in control conditions, and upon the loss of ARID1A, the global enhancer occupancy of ZIC2 was attenuated (Supplementary Fig. 4C & 4D). Specifically, enhancers for essential mesenchymal and CNCC factors, *CDH2* (N-cadherin) and *GLI2* were decreased in both ARID1A and ZIC2 binding in the PT line of the isogenic pair (Fig. 5D).

We further examined ZIC2 genome-wide occupancy and identified ∼5,500 ZIC2 peaks exclusive to the PT lines. These sites did not exhibit ARID1A binding even in CTRL conditions, indicating that *ARID1A*- haploinsufficiency triggered genomic relocation of ZIC2 to non-BAF regulated sites that displayed aberrant accessibility (Fig. 5E; Supplementary Fig. 4E, F). Notably, ZIC2 was bound at regulatory regions for ∼21% of all the differentially upregulated genes in PT lines. Genes associated with aberrantly bound ZIC2 enhancers (Fig. 5F) were enriched for neuronal pathways (Fig. 5G), while ZIC2-bound promoters were associated with common cellular processes (Fig. 5H). These findings were consistent with the upregulation of neuronal genes observed in our *ARID1A*-haploinsufficient PT lines. Collectively, these data suggest that ARID1A-BAF opened the chromatin at EMT enhancers during CNCC specification, allowing for ZIC2 binding. Furthermore, in the absence of ARID1A, ZIC2 relocated to neuronal enhancers whose accessibility was likely modulated by other chromatin remodelers.

### Zic2 is required for NCC EMT and delamination in vivo

To further investigate the role of ZIC2 in EMT and NCC formation *in vitro*, we characterized the spatiotemporal expression of Zic2 during murine neurulation using a previously described *Zic2* reporter mouse line, Tg(*Zic2*^EGFP^)^67^. At E8.0, prior to CNCC delamination and migration (similar to the pre-migratory stage *in vitro*), Zic2 was expressed all along the dorsal neural tube in cells positive for Foxd3, a marker of pre-migratory NCCs (Fig. 6A). By E9.5, there were few Zic2^+^ pre-migratory cells remaining in the dorsal region of the neural tube, indicating successful NCC delamination and migration (Fig. 6B). Additionally, we observed migratory NCCs (Sox10^+^/EGFP) migrating throughout the dorsolateral and ventromedial paths in the mesenchyme (Fig. 6B). This observation suggested that Sox10/EGFP^+^ cells migrating in the mesenchyme (Fig. 6B) were previously Zic2^+^ pre-migratory cells originating from the dorsal region of the neural tube (Fig. 6A).

**Figure 6.**
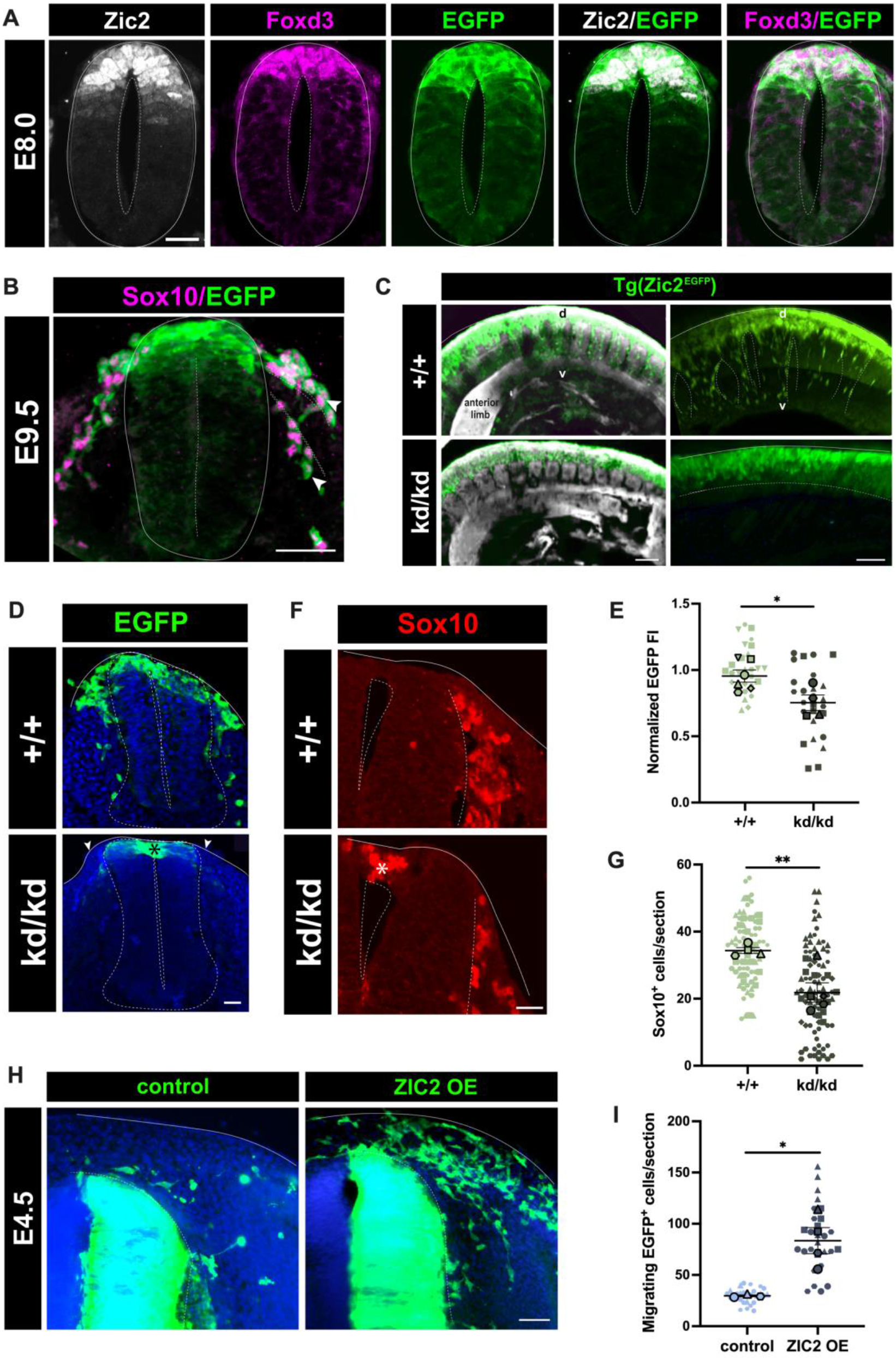
Zic2 activates neural crest EMT and delamination from the neural tube in vivo. (A) A transverse cross-section of the neural tube in an E8.0 Tg(*Zic2*^EGFP^) embryo immunostained for Zic2 and Foxd3. Scale bar = 20µm, 40X magnification (B) A transverse cross-section of the neural tube in an E9.5 Tg(*Zic2*^EGFP^) embryo immunostained for Sox10. White arrowheads highlight Sox10^+^ migrating NCCs are also GFP^+^ depicting migratory NCCs once expressed Zic2 in a pre-migratory setting. Dotted lines show NCCs migrating in the dorsolateral/ventromedial paths. Scale bar = 50µm, 40X magnification. (C) Lateral view of E9.5 whole-mount control [*Zic2*^+/+^;Tg(*Zic2*^EGFP^)] and Zic2 mutant [*Zic2*^kd/kd^;Tg(*Zic2*^EGFP^)] embryos displaying lack of migration into the mesenchyme in Zic2 mutant conditions. Scale bar = 100µm, 5X magnification. d = dorsal; v = ventral. (D) Representative images of a transverse cross-sections of the neural tube from E9.0 control [*Zic2*^+/+^; Tg(*Zic2*^EGFP^)] and Zic2 mutant [*Zic2*^kd/kd^;Tg(*Zic2*^EGFP^)] embryos. Arrowheads highlight the absence of GFP^+^ cells in the dorsal sub-epidermal area of a Zic2-mutant embryo, with GFP^+^ cells accumulating inside the dorsal neural tube represented by *. Scale bar = 10µm, 40X magnification. (E) Superplot quantification of the fluorescent intensity (FI) of GFP^+^ cells in the dorsal sub-epidermis of E9.0 [*Zic2*^kd/kd^; Tg(*Zic2*^EGFP^)] embryos normalized to the control intensity. n = 4-6 independent experiments represented by a distinctive shape with each small data point representing at least four sections per embryo and 4-6 embryos per genotype. The larger data points correspond to the average values of each replicate. A two-tailed unpaired t-test performed on the average values between Zic2-WT and Zic2-mutant embryos; p < 0.05 was considered significant; *p = 0.0255. (F) Representative images of a transverse cross-sections of the neural tube from E10.5 control [*Zic2*^+/+^;Tg(*Zic2*^EGFP^)] and Zic2 mutant [*Zic2*^kd/kd^;Tg(*Zic2*^EGFP^)] embryos immunostained for Sox10. The * highlights an aberrant accumulation of Sox10^+^ cells inside the neural tube at this developmental stage. Scale bar = 20µm, 40X magnification. (G) Superplot quantification of the number of Sox10^+^ cells per section. n = 4-5 independent experiments represented by a distinctive shape with each small data point representing at least five sections per embryo and 4-5 embryos per genotype. The larger data points correspond to the average values of each replicate. A two-tailed unpaired t-test performed on the average values between Zic2-WT and Zic2-mutant embryos; p < 0.05 was considered significant; **p < 0.0074. (H) Transverse cross-sections of the developing neural tube at E4.5 (HH24-25) in chick embryos electroporated at HH9-11 with plasmids carrying Zic2-GFP or GFP alone. OE = over expression. Scale bar = 50µm, 10X magnification (I) Superplot quantification of the number of migrating GFP^+^ cells in sections of E4.5 (HH24-25) chick embryos electroporated at HH9-11 with plasmids carrying ZIC2-GFP or GFP alone. n = 3-4 independent experiments represented by a distinctive shape with each small data point representing at least five sections per embryo and 3-4 embryos per genotype. The larger data points correspond to the average values of each replicate. A two-tailed unpaired t-test performed on the average values between control-GFP and Zic2-GFP; p < 0.05 was considered significant; *p < 0.0161.

In order to assess the impact of Zic2 loss-of-function in NCCs, we crossbred the Tg(*Zic2*^EGFP^) strain with the previously reported *Zic2*^kd^ line^63^ to obtain mutant [*Zic2*^kd/kd^;Tg(*Zic2*^EGFP^)] and control [*Zic2*^+/+^;Tg(*Zic2*^EGFP^)] embryos in the same litter. In control embryos, many EGFP-labeled cells exhibited a stereotypical migration pattern through the dorsal sub-epidermal region of the mesenchyme (Fig. 6C-E). However, in the Zic2 mutant embryos, very few migrating EGFP-labeled cells were detected (Fig. 6C-E). Instead, an aberrant accumulation of EGFP^+^ cells was observed in the dorsal neural tube in the mutants compared to the controls (Fig. 6D). Notably, there was a significant reduction of migratory Sox10^+^ cells detected in the Zic2 mutant embryos (Fig. 6F & 6G). The Sox10^+^ cells were mostly restricted to the dorsal neural tube in the *Zic2*-mutant embryos indicating a lack of EMT, delamination, and migration in the absence of Zic2 (Fig. 6F). Consistent with our *in vitro* data, these results indicate that NCCs failed to delaminate from the neural tube in Zic2 mutant embryos, demonstrating a crucial role for Zic2 in cell delamination via EMT.

### ZIC2 gain-of-function at pre-migratory stages elicits aberrant NCC delamination in vivo

To explore whether Zic2 was sufficient to trigger cell detachment through EMT *in vivo*, we conducted gain-of-function experiments by ectopically expressing human ZIC2 and/or GFP into the neural tube of HH9-11 chick embryos (pre-neurulation). As expected, GFP^+^ cells were largely observed in the neural tube at developmental stage HH24-25 (E4.5; Fig. 6H). Notably, ectopic expression of ZIC2 at the same pre-migratory stages triggered a significantly increased number of cells delaminating from the neural tube (Fig. 6H & 6I). This indicates that ectopic expression of ZIC2 was sufficient to induce delamination in NCCs. Collectively our *in vivo* gain-of-function data further support a key role for ZIC2 in EMT and delamination of NCCs from the neural tube.

## Discussion

ARID1A is one of two integral DNA-binding subunits of the canonical BAF chromatin remodeling complex. While ARID1A-BAF has a wide-range of roles including regulating aberrant chromatin accessibility in cancer contexts^6,68^, its function in physiological craniofacial development is underappreciated. Mutations in *ARID1A* are associated with Coffin-Siris Syndrome, a rare developmental disorder predominantly characterized by coarse craniofacial features^31,32^. The hyperplastic craniofacial skeletal features seen in Coffin-Siris Syndrome are derived from the cranial neural crest^31,33,37^.

Herein, we investigated the role of ARID1A-BAF in the molecular regulation of CNCC formation using *ARID1A*^+/-^ patient-derived iPSCs. It is critical to highlight the considerable incidence of somatic mosaicism associated with *ARID1A* mutations^17,18^. While *de novo* germline mutations have been reported, most CSS patients who harbor mutations in ARID1A are typically mosaic^42^. Recently it was shown that full mutations in *ARID1A* result in a much more severe phenotype, often with multiple congenital anomalies, frequently leading to termination of pregnancy or spontaneous demise during or after birth^42^. In a research context, this mosaicism provides us with an isogenic cellular platform in which the wild-type and *ARID1A*-mutant lines share the same genetic background. This offers the advantage of eliminating confounding genomic factors, ensuring all the cellular phenotypes observed are a direct result of *ARID1A*- haploinsufficiency from a clinical setting.

We previously reported an essential ARID1 subunit switch between ARID1A and ARID1B in BAF that occurs at the transition of iPSC-to-neuroectoderm and neuroectoderm-to-CNCC^3^. Notably, subunit switching for distinct regulatory function is not restricted to BAF as this phenomenon occurs in other chromatin remodelers. For example, among the mutually exclusive ATPases in the NuRD complex, CDH3/4/5, each subunit coordinates specific sets of genes necessary for various stages of cortical development^69^. Ultimately, signaling pathways and interactors that regulate these ARID1 subunit switches and expression timing remain unknown. Similarly, recruitment of BAF to its target regions is globally regulated by the AP-1 transcription factor complex^53^. However, this recruitment process could also be guided by cell type-specific transcription factors essential for NCC formation^45,54–56^ as depicted in our motif analysis of ARID1A-bound regions.

In this study, we have unveiled a novel axis for CNCC formation including ZIC2 and ARID1A-BAF (Fig. 7). ZIC2 is a transcription factor of the GLI superfamily, which is known to regulate several aspects of neurodevelopment^70–72^. Variants of *ZIC2* are associated with holoprosencephaly type 5b^65^, a developmental syndrome that shares craniofacial phenotypes^66^ with Coffin-Siris^32,32^. Holoprosencephaly patients harboring a *ZIC2* mutation present with bitemporal narrowing, upslanting palpebral fissures, a short nose with anteverted nares, a broad and well demarcated philtrum, and large ears^66^. Notably, these craniofacial anomalies do not appear in holoprosencephaly patients with mutations in other causative genes^66^.

**Figure 7.**
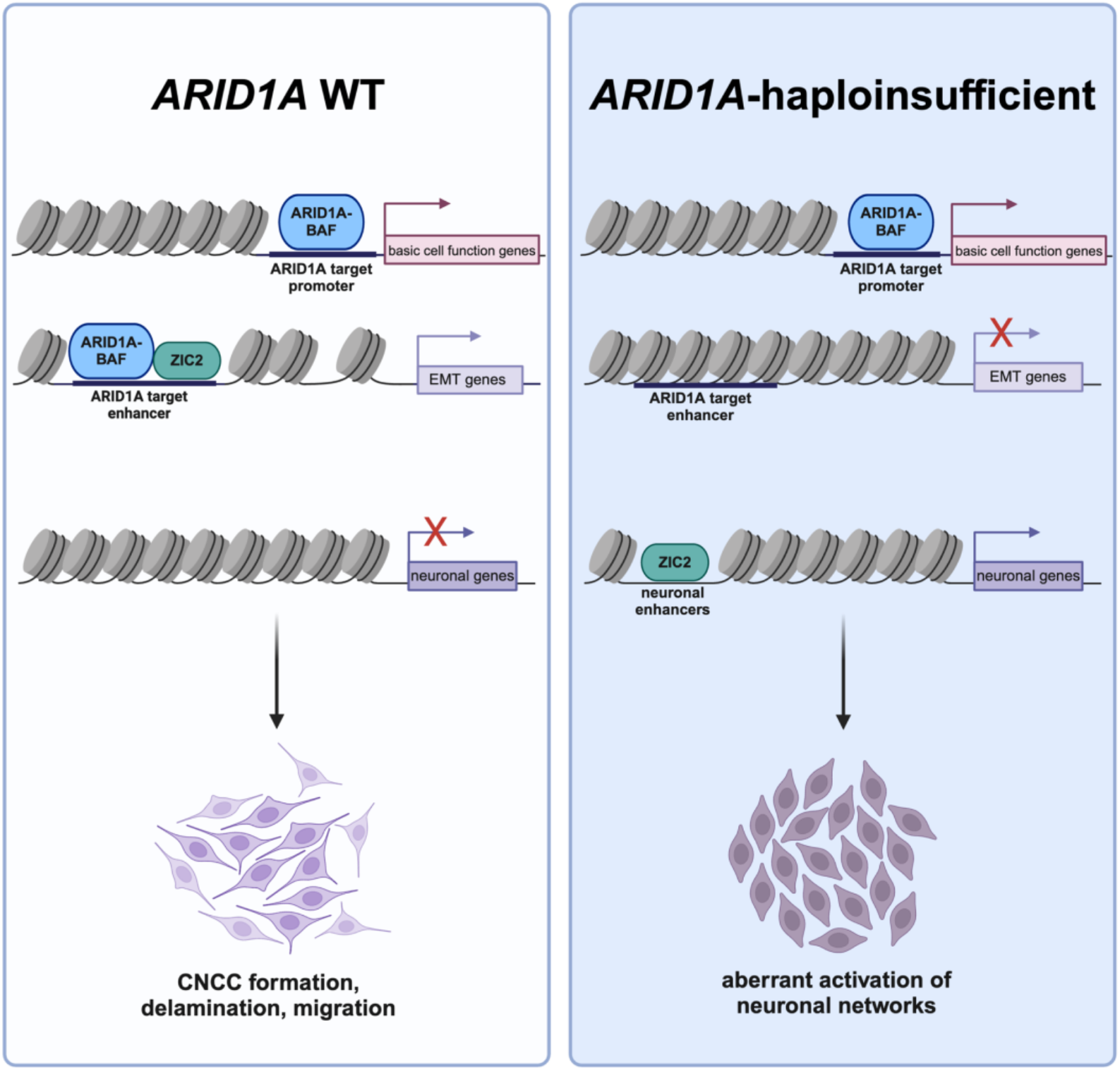
Working model of ARID1A-BAF and ZIC2 regulating EMT networks at enhancers during CNCC specification. In wild-type (WT) ARID1A conditions during CNCC specification, ARID1A-BAF binds at target promoters regulating basic cellular functions. ARID1A-BAF regulates the accessibility of EMT enhancers allowing for ZIC2 to bind and regulate the expression of EMT networks for CNCC specification. In *ARID1A*- haploinsufficient lines, the limited WT ARID1A protein encoded from the single healthy ARID1A allele preferentially binds to ARID1A-target promoters to regulate basic cell functions while binding at enhancers for EMT network activation is dispensable. In the absence of ARID1A-BAF-regulated accessibility at EMT enhancers, ZIC2 relocates to alternate regulatory regions activating the default neuronal networks.

Our *in vitro* and *in vivo* data, encompassing human iPSCs, mouse models, and chick models, support a mechanism in which ARID1A opens the chromatin for ZIC2 at the onset of EMT to induce NCC delamination from the neural tube. With multiple *in vivo* models, we demonstrate that ZIC2 is indispensable for NCC EMT and subsequent migration and discovered the molecular foundation of this regulation through ARID1A-mediated accessibility at EMT enhancers. Supporting this model *in vivo*, Zic2 is expressed in mouse pre-migratory NCCs and is required for their delamination from the neural tube. Conversely, ectopic expression of ZIC2 in chick embryos displays asynchronous delamination and EMT.

In *ARID1A*^+/-^ conditions, accessibility at EMT enhancers is perturbed, preventing lineage-specific transcription factors, like ZIC2, from binding at these elements to promote EMT. In the absence of ARID1A, ZIC2 relocates to neuronal enhancers, suggesting ZIC2 plays a role in the aberrant activation of neuronal networks. As both NCCs and neurons are derived from the neuroectoderm^35,73–76^, and neuroectoderm-like cells are successfully specified in *ARID1A*-haploinsufficiency, our results potentially suggest a default trajectory towards the neuronal lineage when the neural crest lineage cannot be specified. However, further studies will be needed to validate this model.

ZIC2 has multifaceted functions in neuronal development^70–72^, hindbrain patterning^60,77^, and axon guidance^67,78–80^. Zic2 controls axon midline repulsion and this neurodevelopmental process has been recently proposed to be influenced by enhancers containing Zic2 binding motifs^81^. Thus, the genomic relocation of ZIC2 and aberrant upregulation of neuronal-associated genes is consistent with its known role in regulating neuronal processes. However, additional studies will be required to determine the molecular regulation and contribution from other lineage-specific transcription factors and chromatin remodelers underlying this relocation.

Notably, EMT is a main function of other non-developmental processes including wound healing^82^ and fibrosis^83,84^. Therefore, it would be crucial to investigate whether ARID1A-BAF directs EMT in these settings and if distinctive cell type-specific transcription factors are involved. Additionally, EMT extensively modulates cancer progression and metastasis^85^. ARID1A is mutated in ∼10% of all tumors with postulated balancing oncogenic^20,21^ and tumor suppressive functions^22–25,68^. However, ARID1A is predominantly reported to inhibit EMT in a cancer context^86–88^. This may suggest a niche role for ARID1A-BAF in promoting EMT exclusively in embryonic development, but further studies will be needed to support this hypothesis.

In summary, this study provides novel insights into how chromatin remodelers regulate neural crest specification, delamination, and migration, suggesting a potential pathological mechanism underlying severe craniofacial phenotypes in developmental syndromes.

## Materials and Methods

### Human iPSC culture

The CTRL1 iPSC line was obtained from the iPSC Core at the University of Pennsylvania (SV20, male, age 43). Skin fibroblasts were obtained from two pediatric Coffin-Siris patients (PT1 and PT2) by the team of Dr. Gijs Santen at Leiden University. PT1 is a teenage male, while PT2 is a female.

We complied with all relevant ethical regulations for work with human participants and informed consent was obtained by Leiden University (LUMC), which approved the protocol under the coordination of Dr. Gijs Santen. The study was conducted in accordance with the criteria set by the Declaration of Helsinki. The protocol to make iPSCs was approved by the IRB at LUMC. Collecting patient material and establishing iPSCs were all performed according to local (LUMC) IRB protocols.

The fibroblasts were reprogrammed into iPSCs with the polycistronic lentiviral vector LV.RRL.PPT.SF.hOKSM.idTomato.-preFRT^89^ as described elsewhere^90^. Multiple clones per line were derived and one *ARID1A* mutant clone per line was used for this study: PT1 = LUMC0081iARID02 or 81-2 and PT2 = LUMC0064iARID04 or 64-4.

Since PT2 exhibited somatic mosaicism, *ARID1A* wild type iPSC clones were also isolated from PT2 (CTRL2; LUMC0064iARID09 or 64-9). For each clone per line, pluripotency was assessed by immunofluorescence microscopy using antibodies against NANOG, OCT3/4, SSEA4, and Tra-1-81 under maintenance conditions and antibodies against (TUBB3, AFP, and CD31) after spontaneous differentiation into the three germ layers as described elsewhere^89,91^. Clones with proper pluripotent characteristics were selected for downstream usage. Karyotyping by G binding was assessed for all three lines by the Leiden University Medical Center and short tandem repeat profiling was performed by the Leiden University Medical Center and all four lines were profiled at the Stem Cell and Regenerative Neuroscience Center at Thomas Jefferson University. The iPSC lines were expanded in mTeSR1 medium (STEMCELL Technologies) on Matrigel (BD Pharmingen). Cells were passaged ∼1:10 at 80% confluency using 0.5mM EDTA (STEMCELL Technologies) and small cell clusters (50–200 cells) were subsequently plated on tissue culture dishes coated overnight with GeltrexTM LDEV-Free hESC-qualified Reduced Growth Factor Basement Membrane Matrix (Fisher-Scientific).

### RNA extraction and sequencing

Cells were lysed in Tri-reagent (Zymo) and total RNA was extracted using Direct-zol RNA Miniprep kit (Zymo) according to the manufacturer’s instructions. RNA was quantified and RNA integrity was checked by Tapestation 4150 (Agilent). Only samples with RIN value above 8.0 were used for transcriptome analysis. RNA libraries were prepared using 1μg of total RNA input using NEB-Next® Poly(A) mRNA Magnetic Isolation Module, NEBNext UltraTM II Directional RNA Library Prep Kit for Illumina and NEBNext UltraTM II DNA Library Prep Kit for Illumina according to the manufacturer’s instructions (New England Biolabs). Libraries were sequenced on a NextSeq2K (Illumina) generating single-end ∼138 bp reads.

### RNA-sequencing analysis

After removing the adapters with TrimGalore!. Kallisto^92^ was used to quantify reads mapping to each gene. Differential gene expression levels were determined using DESeq2^93^. All statistical analyses were performed using the latest versions of BEDTools^94^, deepTools^95^, R, and GraphPad.

### In vitro immunofluorescence

Immunohistochemistry of iPSCs and CNCCs was performed in µ-Slide 4 Well Glass Bottom (IBIDI 80426). Upon fixation (4% PFA for 10 minutes), cells were permeabilized in blocking solution (0.1% Triton X-100, 1x PBS, 5% normal donkey serum) and then incubated with the antibody of interest. The total number of cells in each field was determined by counterstaining cell nuclei with 4,6-diamidine-2-phenylindole dihydrochloride (DAPI; Sigma-Aldrich; 50 mg/ml in PBS for 15 min at RT). Immunostained cells were analyzed via confocal microscopy using a Nikon A1R+. Images were captured with a 4x objective for iPSCs and a 60x objective for CNCCs with a pinhole of 1.0 Airy unit. Analyses were performed in sequential scanning mode to rule out cross-bleeding between channels. Fluorescence intensity and counting quantifications were performed with Fiji and the NIS-Elements AR software. In detail, to analyze the net intensity of the cells, we create a mask around each nucleus using DAPI intensity as the base criterion. This mask served as the reference for measuring the fluorescence intensity or number of cells positive for nuclear proteins. To analyze cytoplasmic proteins, the previously drawn nucleus mask was expanded to encompass the entire cytoplasm. All antibodies are listed in Supplementary File 6. All two-tailed unpaired t-tests were performed in GraphPad.

### CNCC specification

The iPSC lines were specified into CNCC as previously described^3,45^. In brief, iPSCs were cultured with CNCC specification media: 1:1 Neurobasal medium/D-MEM F-12 medium (Invitrogen), 1X penicillin-streptomycin solution, 1x Glutamax supplement (100x stock, Invitrogen), 0.5x B-27 supplement with Vitamin A (50x stock, Invitrogen), 0.5x N-2 supplement (100x stock, Invitrogen), 5 μg/ml bovine insulin (Sigma-Aldrich), 20 ng/ml EGF (Sigma-Aldrich), and 20 ng/ml hFGF (Biolegend) for 6 days. Medium was changed every 1-2 days. At day ∼7 when early migratory CNCCs first appeared, the cells were transitioned to CNCC early maintenance media: 1:1 Neurobasal medium/D-MEM F-12 medium (Invitrogen), 1X penicillin-streptomycin solution, 1x Glutamax supplement (100x stock, Invitrogen), 0.5x B-27 supplement with Vitamin A (50x stock, Invitrogen), 0.5x N-2 supplement (100x stock, Invitrogen), 1 mg/ml bovine serum albumin, serum replacement grade (Gemini Bio-Products # 700-104 P), 20 ng/ml EGF (Sigma-Aldrich), and 20 ng/ml hFGF (Biolegend) for 7 days. After 14 full days of CNCC specification, the cells were maintained in an “early CNCC” state for up to 21 days for subsequent experiments, or immediately transitioned to CNCC late maintenance media: 1:1 Neurobasal medium/D-MEM F-12 medium (Invitrogen), 1X penicillin-streptomycin solution, 1x Glutamax supplement (100x stock, Invitrogen), 0.5x B-27 supplement with Vitamin A (50x stock, Invitrogen), 0.5x N-2 supplement (100x stock, Invitrogen), 1 mg/ml bovine serum albumin, serum replacement grade (Gemini Bio-Products # 700-104 P), 20 ng/ml EGF (Sigma-Aldrich), 20 ng/ml hFGF (Biolegend), 3µM ChIRON 99021 (Selleck Chem S1263), and 50ng/mL BMP2 (Peprotech 120-02). CNCCs transitioned to late maintenance media can be maintained for up to 2 additional weeks.

### Flow cytometry

CTRL and PT iPSCs were treated with accutase for 5 minutes to obtain a single-cell suspension. Cells were then washed with cold 1X phosphate buffered saline (PBS) + 2% fetal bovine serum (FBS) and live cells were counted and resuspended in 1X PBS to 1x10^6^ cells/mL. LIVE/DEAD aqua stain (ThermoFisher L34965) was prepared following the manufacturer’s instructions and 1ul of prepared stain was added per 1x10^6^ cells and incubated 30 min at 4°C in the dark. Cells were then washed twice in 1X PBS + 2% FBS. Cells were resuspended in up to 100ul 1X PBS + 2% FBS and 2ul of respective antibodies (1:50 dilution; Supplementary File 6) and incubated 15 min at 4°C in the dark. The antibodies were removed, and the cells were washed once in 1X PBS + 2% FBS. Stained cells were resuspended in 300ul of PBS + 2% FBS and strained using a 35µM strainer. Flow cytometry data were acquired using a BD Celesta and analyzed with FlowJo Software v10.10.

### Western Blotting

Cells were washed three times in 1X PBS and lysed in radioimmunoprecipitation assay buffer (RIPA buffer; 50 mM Tris-HCl pH7.5, 150 mM NaCl, 1% Igepal, 0.5% sodium deoxycholate, 0.1% SDS 500 μM DTT) with protease inhibitors. Between 20-30 μg of whole cell lysate were loaded in Novex WedgeWell 4–20% Tris-Glycine Gel (Invitrogen) and subject to SDS-PAGE. Proteins were then transferred to a Immun-Blot PVDF or Nitrocellulose membrane (ThermoFisher) for antibody probing. Membranes were blocked with a 10% BSA in TBST solution for 30 minutes then incubated with diluted primary antibodies (Supplementary File 6) in 5% BSA in TBST solution. Membranes were then washed with TBST and incubated with diluted secondary antibodies (Supplementary File 6). Chemiluminescent signal was detected using the KwikQuant Western Blot Detection Kit (Kindle BioSciences) or the Amersham ECL Prime Western Blotting Detection Reagents (Cytiva) and a KwikQuant Imager.

### Transwell

The transwell assay was performed using the Cell Migration Assay Kit (abcam ab235694) following the manufacturer’s protocol. Briefly, CNCCs were cultured to ∼80% confluency and growth factors (EGF and FGF) were removed from the CNCC early maintenance media for 24 hours. The bottom chamber of the migration plate was prepared with growth factor-free CNCC early media and 10% FBS and the top chamber was assembled. Cells were harvested and counted, and 200,000 cells were resuspended in growth factor-free CNCC early maintenance media and seeded in the top chamber. Cells were incubated in the migration chamber at 37°C for 48 hours. After 48 hours, the cells that invaded through the migration membrane were dissociated and incubated in Cell Dye for 1 hour at 37°C. One set of standards per cell type was created via serial dilution and incubated in Cell Dye for 1 hour at 37°C. Absorbance for each sample and standard was read at Ex/Em-530/590nm using a PolarStar Optima plate reader (BMG LabTech). Percent migration was calculated from the linear curve of the plotted standards.

### Scratch wound assay

CNCCs were cultured to ∼80% confluency and growth factors (EGF and FGF) were removed from the CNCC early maintenance media for 24 hours. Cells were harvested and counted, and 100,000 cells were resuspended in growth factor-free CNCC early maintenance media and seeded in an IncuCyte ImageLock 96-well plate (Sartorius 4379) in triplicate and incubated for 48 hours at 37°C. The IncuCyte 96-Well Woundmaker Tool was used to create identical wounds in each well. The wells were washed twice to remove the cells dislodged from the Woundmaker Tool and the media was refreshed. The cells were incubated at 37°C, and migration was assayed for 96 hours with repeat scanning every 2 hours via the IncuCyte ZOOM Live-Cell Analysis System.

### ChIP-sequencing

All samples from different conditions were processed together to prevent batch effects. Approximately 13 million cells were cross-linked with 1% formaldehyde for 5 minutes at room temperature, quenched with 125 mM glycine, harvested, and washed twice with 1X PBS. The fixed cell pellet was resuspended in ChIP lysis buffer (150 mM NaCl, 1% Triton X-100, 0.7% SDS, 500 μM DTT, 10 mM Tris-HCl, 5 mM EDTA) and chromatin was sheared to an average length of 200–900 base-pairs, using a Covaris S220 Ultrasonicator at 5% duty factor between 6-9 minutes. The chromatin lysate was diluted with SDS-free ChIP lysis buffer. 15μg of antibody was used for ARID1A and ZIC2 and 3μg of antibody for H3K27ac (Supplementary File 6). The antibody was added to at least 5μg of sonicated chromatin along with Dynabeads Protein G magnetic beads (Invitrogen) and incubated with rotation at 4°C overnight. The beads were washed twice with each of the following buffers: Mixed Micelle Buffer (150 mM NaCl, 1% Triton X- 100, 0.2% SDS, 20 mM Tris-HCl, 5 mM EDTA, 65% sucrose), Buffer 200 (200 mM NaCl, 1% Triton X-100, 0.1% sodium deoxycholate, 25 mM HEPES, 10 mM Tris-HCl, 1 mM EDTA), LiCl detergent wash (250 mM LiCl, 0.5% sodium deoxycholate, 0.5% NP-40, 10 mM Tris-HCl, 1 mM EDTA) and a final wash was performed with cold 0.1X TE. Finally, beads were resuspended in 1X TE containing 1% SDS and incubated at 65°C for 10 min to elute immunocomplexes. The elution was repeated twice, and the samples were incubated overnight at 65°C to reverse cross-linking, along with the input (5% of the starting material). The DNA was digested with 0.5 mg/ml Proteinase K for 1 hour at 65°C and then purified using the ChIP DNA Clean & Concentrator kit (Zymo) and quantified with QUBIT. Barcoded libraries were made with NEBNext Ultra II DNA Library Prep Kit for Illumina) using NEBNext Multiplex Oligos Dual Index Primers for Illumina (New England BioLabs) and sequenced on NextSeq2K (Illumina) producing ∼138 bp single-end reads.

### ChIP-seq analysis

After removing the adapters with TrimGalore!, the sequences were aligned to the reference hg19, using Burrows-Wheeler Alignment tool, with the MEM algorithm^96^. Uniquely mapping aligned reads were filtered based on mapping quality (MAPQ > 10) to restrict our analysis to higher quality and uniquely mapped reads, and PCR duplicates were removed. MACS2^97^ was used to call peaks using the default parameters at 5% FDR. All statistical analyses were performed using the latest versions of BEDTools^94^, deepTools^95^, R, and GraphPad.

### ATAC-sequencing

For ATAC-seq experiments, 50,000 cells per condition were processed as previously described^98^. Briefly, 50,000 cells were collected, washed, and lysed. The chromatin was subjected to transposition/library preparation via a Tn5 transposase using the Tagment DNA Enzyme and Buffer Kit (Ilumina 20034197) and incubated at 37°C for 30 min with slight agitation. Transposed DNA was purified using a MinElute PCR Purification Kit (Qiagen 28004). Transposed DNA fragments were then amplified using a universal and barcoded primer^98^. Thermal cycling parameters were set as follows: 1 cycle of 72°C for 5 min, 98 °C for 30 seconds, followed by 5 cycles of 98°C for 10 seconds, 63°C for 30 seconds, and 72°C for 1 min. The amplification was paused and 5 µl of the partially amplified, transposed DNA was used for a qPCR side reaction including the universal and sample-specific barcoded primers^98^, PowerUp SYBR Green Master Mix (Applied Biosystems), NEBNext High-Fidelity 2x PCR Master Mix, and nuclease-free water. The qPCR side reaction parameters were set as follows: 1 cycle of 72°C for 5 min, 98°C for 30 seconds, followed by 40 cycles of 98°C for 10 seconds, 63°C for 30 seconds, and 72°C for 1 min. The Rn vs cycle was plotted to determine the remaining number of PCR cycles needed where 1/3 of the maximum fluorescent intensity corresponds to the additional cycle number. The remaining partially amplified transposed DNA was fully amplified using the previous parameters with the additional cycle number determined from the qPCR side reaction. The amplified, transposed DNA was purified using AMPure XP beads (Beckman Coulter A63881) and sequenced using an Illumina NextSeq2K, generating ∼138 bp single-end reads.

### ATAC-sequencing analysis

After removing the adapters with TrimGalore!, the sequences were aligned to the human reference genome, hg19, using the Burrows–Wheeler Alignment tool, with the MEM algorithm^96^. Aligned reads were filtered based on mapping quality (MAPQ > 10) to restrict our analysis to higher quality and uniquely mapped reads, and PCR duplicates were removed. All mapped reads were offset by +4 bp for the forward strand and -5 bp for the reverse strand. MACS2^97^ was used to call peaks using the default parameters at 5% FDR. All statistical analyses were performed using the latest versions of BEDTools^94^, deepTools^95^, R, and GraphPad.

## Motif analysis

Fasta files for the regions of interest were produced using BEDTools^94^. Motif analysis of all lost ARID1A- bound regions and ARID1A-bound promoters was performed using HOMER^99^. Shuffled input sequences were used as background. E-values < 0.001 were used as a threshold for significance. Transcription factor motif quantification was performed using FIMO^100^ (The MEME Suite) on a background of lost ARID1A regions.

### Mouse lines

The Tg(*Zic2*^EGFP^) HT146Gsat/Mmcd line (identification number RP23-158G6) was generated by GENSAT^101^ and obtained from the Mutant Mouse Regional Resource Center (http://www.mmrrc.org/strains/17260/017260.html). The hypomorphic *Zic2*^kd/kd^ mouse line (*Zic2^tm1Jaru^*;MGI:2156825) was obtained from the RIKEN repository. *Zic2*^+/+^;Tg(*Zic2*^EGFP^) and *Zic2*^kd/kd^;Tg(*Zic2*^EGFP^) embryos were obtained from breeding *Zic2*^+/kd^;Tg(*Zic2*^EGFP^) mice. The day a vaginal plug appeared was considered embryonic day E0.5. All mouse lines were congenic on a C57BL/6J background and were kept in a timed pregnancy-breeding colony at the Instituto de Neurociencias. The animal protocols were approved by the Institutional Animal Care and Use Committee and met European and Spanish regulations.

### Constructs and in ovo electroporation

Fertilized White Leghorn chicken eggs were incubated at 38°C until the desired developmental stage according to Hamburger and Hamilton^102^. Platinum electrodes were used with 5 x 10 ms pulses at 25 V generated with a TSS20 Ovodyne electroporator. The following plasmids were injected at the indicated concentrations: pCAG-ZIC2 plasmid (1 µg/µl) and CAG-GFP (0.5 µg/µl). Due to technicalities underlying electroporation, all analyses of electroporated chicken embryos were confined to the trunk region.

### In vivo immunohistochemistry and iDISCO

For immunohistochemistry (IHC), chick embryos fixed overnight at 4°C, and mouse embryos fixed for 4- 5hrs (E8.5-E9.5) with 4% paraformaldehyde (PFA)/phosphate-buffered saline (PBS) and were washed with PBS. Cryostat (40µm) and vibratome (70µm) sections or whole embryos were incubated with the corresponding primary and secondary antibodies according to standard protocols. For iDISCO, embryos were fixed overnight at 4°C with 4% PFA/PBS, and immunolabeling was performed before clarification.

The iDISCO protocol was performed as previously published^103^. IHC images were captured with an Olympus FV1000 confocal IX81 microscope/FV10-ASW Software. Images of iDISCO-clarified embryos were obtained with an Ultramicroscope II (LaVisionBiotec). All antibodies are listed in Supplementary File 6.

### In vivo immunohistochemistry analysis

The number of Sox10^+^ or FoxD3^+^ cells per slice was quantified and normalized by neural tube length and the ratio of positive cells inside the tube compared with the total number of cells in each cryosection (30µm). A region of interest was delineated in the migration zone outside the neural tube, and the fluorescence intensity was measured (IntDen parameter in Fiji/ImageJ) and normalized to total fluorescence. Two complementary regions of interest were delineated along the migratory stream (one in the ventromedial area and another in the dorsolateral area). GFP fluorescence intensity was measured, and the total fluorescence area was calculated as: integrated density (fluorescence of the selected area - mean background fluorescence) in Fiji/ImageJ. All two-tailed unpaired t-tests performed in GraphPad.

### Statistical and genomic analysis

All statistical analyses were performed using the latest versions of BEDTools^94^, deepTools^95^, R, and GraphPad. Superplot quantification represent at least n = 3 represented by a distinctive shape with each small data point representing a captured image. The larger data points correspond to the average values of each replicate. A two-tailed unpaired t-test performed on the average values. A value of p < 0.05 was considered significant; *p < 0.05; **p < 0.01; ***p < 0.001; ****p<0.0001; n.s., not significant. Pathway analysis was performed with Ingenuity-Pathway Analysis Suite (QIAGEN Inc., https://digitalinsights.qiagen.com/IPA).

## Acknowledgements

The authors are grateful to the families of the CSS patients for donating the samples, and to Sandra Deliard and Alessandro Gardini for the use of the IPA software and for insightful comments. The authors thank the Genomic Facility at The Wistar Institute (Philadelphia, PA) for the Next Generation Illumina Sequencing. For this work, MT was funded by the G. Harold and Leila Y. Mathers Foundation.

## Data availability

The original genome-wide data generated in this study have been deposited in the GEO database under accession code GSE261846.

## Author contributions

MT, EH, and SMB designed the project. SMB performed all the human iPSC and CNCC experiments and analyzed all the genomic data. AGG, CMP, and MTLC performed all the *in vivo* experiments. CS performed all the iPSC and CNCC immunofluorescence and contributed to data analysis. FJW, MEC, and JK contributed to some of the experiments. GWES recruited the patients and obtained the skin fibroblasts. HMMM reprogrammed the patient fibroblasts into iPSCs and performed quality checks. KAP, DT, SBM and SAB contributed to data analysis and offered critical support. SMB and MT analyzed the data and wrote the manuscript. All the authors read and approved the manuscript.

