## Supplementary Figures for "ARID1A-BAF coordinates ZIC2 genomic occupancy for epithelial to mesenchymal transition in cranial neural crest lineage commitment"

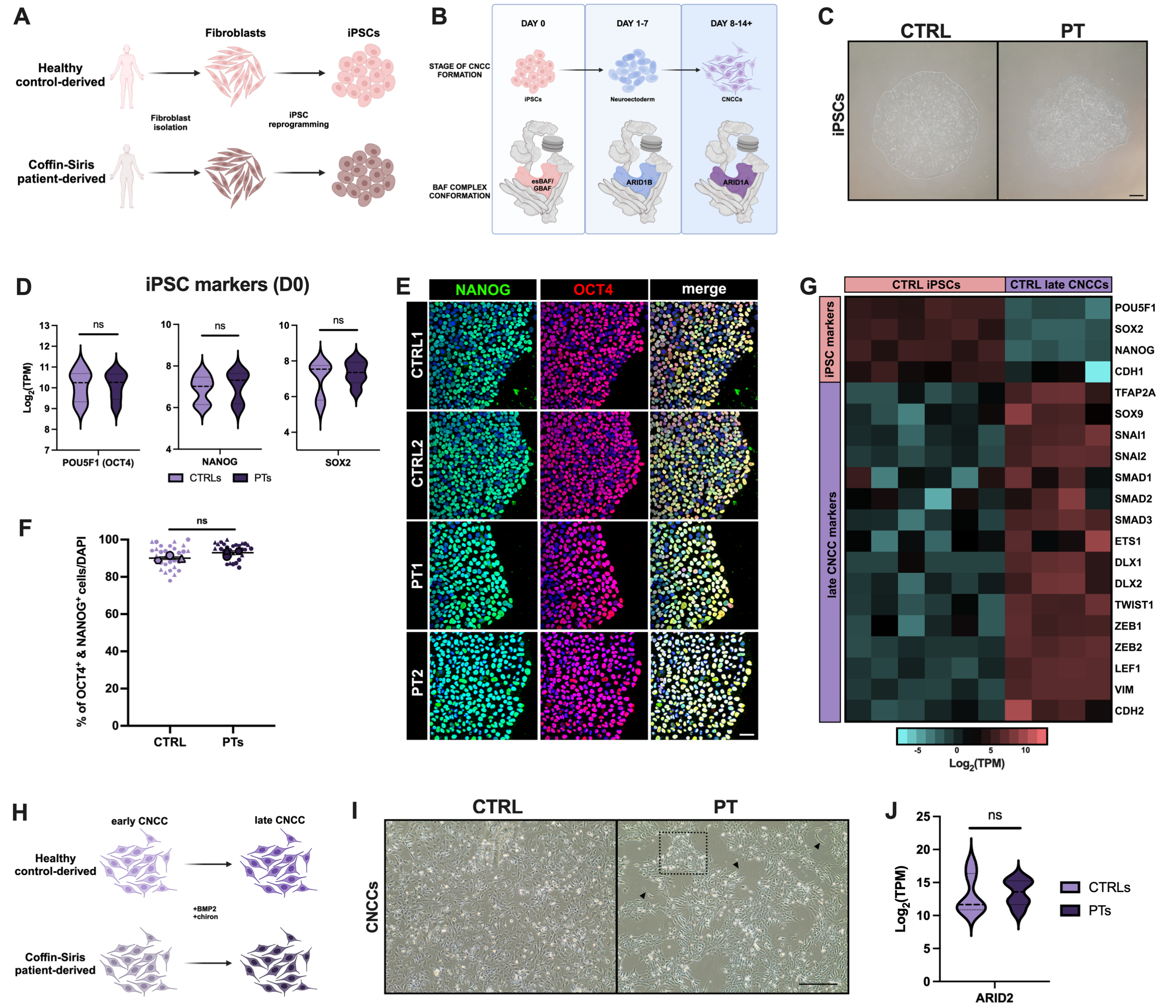


**Supplemental Figure 1 – Validation of iPSC reprogramming and *in vitro* CNCC specification in CTRL and PT cell lines.** (A) Graphical illustration of iPSC reprogramming from ARID1A-haploinsufficient Coffin Siris patient-derived tissue and healthy control tissue. Made with BioRender.com. (B) Visual schematic of ARID1 subunit switching at different morphological stages throughout CNCC specification. Made with BioRender.com. (C) Representative brightfield images of CTRL and PT iPSC colonies show no difference in cellular morphology. Scale bar = 300µm, 4X magnification. (D) Violin plots displaying $\log_2$(TPM) of iPSC markers, POU5F1 (OCT4), NANOG, and SOX2 in CTRL and PT iPSCs. There is no significant difference in expression of pluripotent factors between CTRLs and PTs. A two-tailed unpaired t-test was performed and $p < 0.05$ was considered significant; POU5F1 (OCT4): ns $p = 0.992$, NANOG: ns $p = 0.7227$, and SOX2: ns $p = 0.5268$. (E) Representative images and (F) quantification of an immunofluorescence for pluripotency factors OCT4 and NANOG performed in CTRL and PT iPSCs. (E) DAPI staining on nuclei in blue. Images shown are taken at 20X magnification; scale bar = 50µm. (F) SuperPlot quantification of percentage of OCT4/NANOG double-positive cells per DAPI. n = 3 represented by a distinctive shape with each small data point representing a captured image. The larger data points correspond to the average values of each replicate. A two-tailed unpaired t-test performed on the average values between CTRLs and PTs; $p < 0.05$ was considered significant; ns $p = 0.0894$. (G) Heatmap of the expression of pluripotent markers and CNCC markers in CTRL iPSCs (D0) and late specified CNCCs (+BMP2 and CHIR-99021). Pluripotent genes, including

OCT4, are deactivated while CNCC specifiers are upregulated in CNCCs relative to iPSCs. iPSC columns represent 3 individual replicates for CTRL1 and CTRL2 and the late CNCC columns represent 2 individual replicates from 2 separate CNCC specifications for CTRL1 and CTRL2. (H) Schematic of "early" to "late" CNCC specification through the addition of exogenous BMP2 and CHIR-99021 (GSK3 inhibitor). Made with BioRender.com. (I) Representative brightfield images of D14 CNCCs demonstrating altered morphology of PT CNCCs compared to CTRLs. Black arrows point to examples of aberrant elongated morphology. Black dotted box highlights an example of aberrant cell clustering. Scale bar = 300µm, 10X magnification. (J) Violin plots displaying $\log_2$(TPM) expression of ARID2 at D10 of CNCC specification. There is no significant difference in expression between CTRLs and PTs. A two-tailed unpaired t-test was performed and $p < 0.05$ was considered significant; $p = 0.8420$.

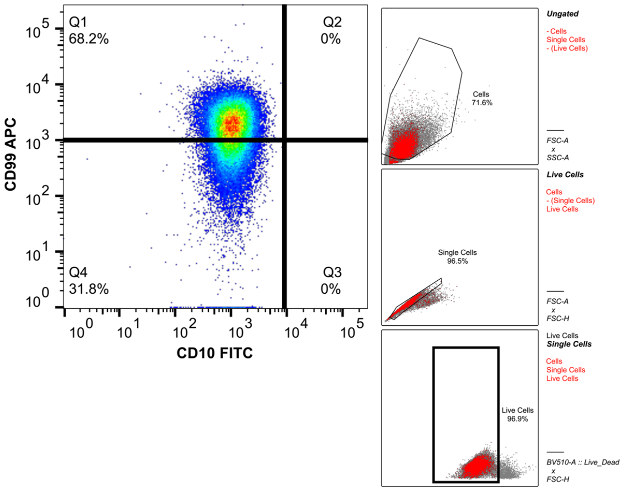


**Supplemental Figure 2 – Flow cytometry back-gating analysis for Figure 1C**. Forward scatter area (FSC-A) versus side scatter area (SCC-A) was used to select the population of cells. FSC-A versus forward scatter height (FSC-H) was used to select single cells and exclude doublets. Subsequent gating was used to identify stained live cells (BV510-A vs FSC-H). CNCC-positive populations were characterized by CD99-APC versus CD10-FITC surface marker expression.

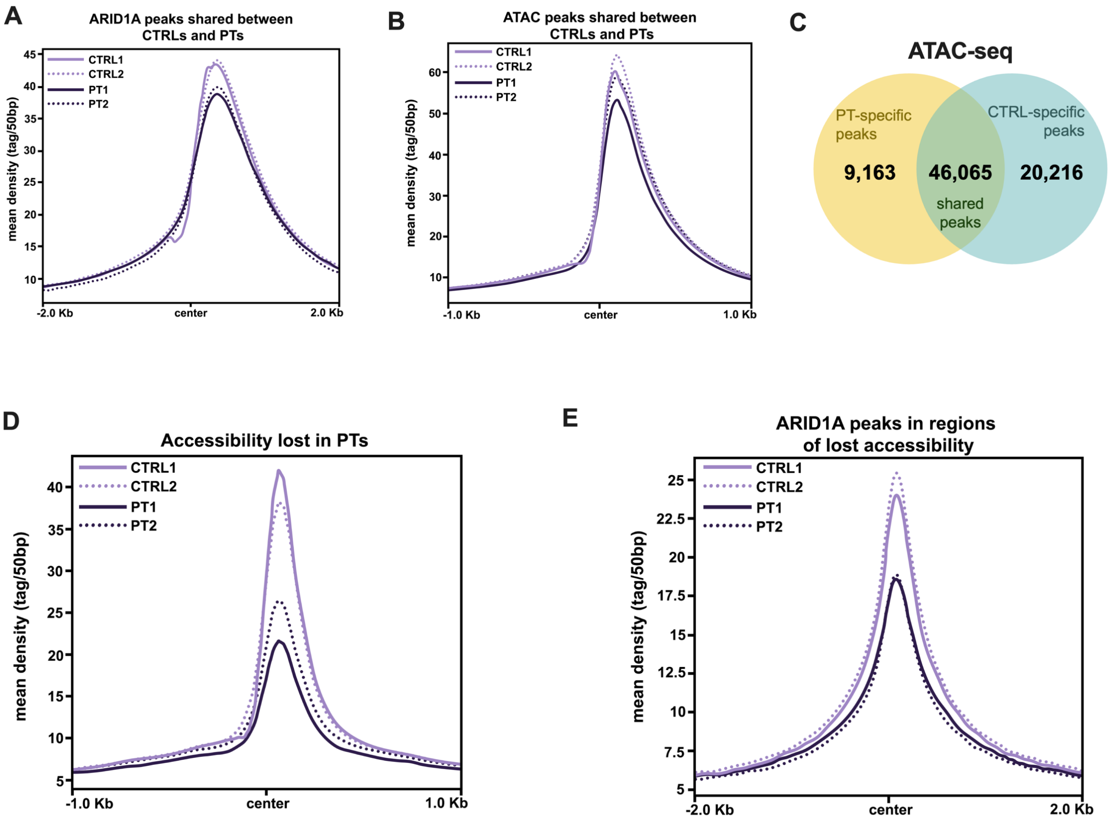


**Supplemental Figure 3 – Global ARID1A binding and accessibility across CTRL and PT D10 CNCCs.** (A) Average profile of ARID1A peaks shared between CTRLs and PTs at D10 of CNCC specification (n = 21,724). Center represents the average overlapping shared binding of ARID1A across the genome. (B) Average profile of regions of accessibility (ATAC peaks) conserved between CTRLs and PTs at D10 of CNCC specification (n = 46,065). Center represents the average overlapping shared ATAC peaks across the genome. C) Venn diagram displaying the number of PT-specific ATAC peaks (ATAC peaks "gained" in PTs; 9,163), CTRL-specific ATAC peaks (ATAC peaks "lost" in PTs; 20,216), and shared ATAC peaks between CTRLs and PTs (conserved regions of accessibility; 46,065) via an ATAC-seq performed at D10 of CNCC specification. (D) Average profile of ATAC peaks lost in PTs at D10 of CNCC specification (n = 20,216). Center represents the average overlapping lost ATAC peaks across the genome. (E) Average profile of ARID1A peaks at regions of lost accessibility in the PTs at D10 of CNCC specification (n = 20,216). Center represents the average overlapping binding of ARID1A at regions of lost accessibility in PTs across the genome.

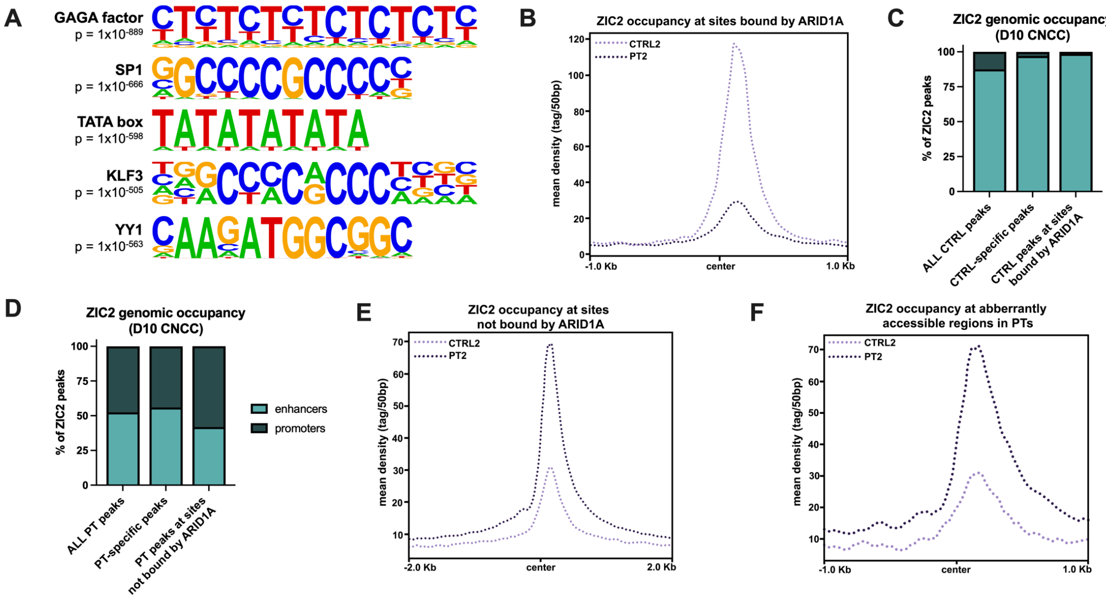


**Supplementary Figure 4 – Genomic features of ARID1A-bound promoters and ZIC2 binding at D10 of CNCC specification.** (A) Motif analysis at ARID1A-bound promoters identified by HOMER. (B) Average profile of ZIC2 peaks at ARID1A sites at D10 of CNCC specification in the isogenic system (n = 358). Center represents the average overlapping binding of ZIC2 at ARID1A-bound regions across the genome. (C) Stacked bar plot depicting the percentage of ZIC2 peaks enriched at enhancers (>1kb from the closest transcription start site or TSS) and promoters (<1kb from the closest TSS) at all CTRL ZIC2 peaks (including peaks shared with PT lines; 87% at enhancers, 13% at promoters), CTRL-specific peaks (ZIC2 peaks exclusive to the CTRLs; 97% at enhancers, 3% at promoters), and overlapping ZIC2- and ARID1A-bound sites in CTRLs (99% at enhancers, 1% at promoters). (D) Stacked bar plot depicting the percentage of ZIC2 peaks enriched at enhancers (>1kb from the closest transcription start site or TSS) and promoters (<1kb from the closest TSS) at all PT ZIC2 peaks (including peaks shared with CTRL lines; 52% at enhancers, 48% at promoters), PT-specific peaks (ZIC2 peaks exclusive to the PT; 56% at enhancers, 44% at promoters), and ZIC2 binding in the PT at regions not bound by ARID1A (42% at enhancers, 58% at promoters). (E) Average profile of ZIC2 peaks at sites not bound by ARID1A at D10 of CNCC specification in the isogenic system (n = 5,460). Center represents the average overlapping binding of ZIC2 at non-ARID1A bound regions across the genome. (F) Average profile of ZIC2 peaks at aberrantly accessible regions in PT2 at D10 of CNCC specification (n = 244). Center represents the average overlapping binding of ZIC2 at PT-specific ATAC peaks across the genome.
